# Phylogenetic Analysis and Structural Evaluation of *Staphylococcus aureus* Serine-Aspartate Repeat-Containing Protein D with a Focus on Periprosthetic Joint Infection

**DOI:** 10.64898/2026.05.01.722179

**Authors:** Kemin Tan, Kathleen A. O’Connor, Xiaobing Zuo, Priyanka Gade, Elijah J. Garcia, Angela Tan, Adam K. Nijhawan, Mike Endres, Youngchang Kim, Kerryl E. Greenwood-Quaintance, Robin Patel, Andrzej Joachimiak

## Abstract

Serine–aspartate repeat-containing protein D (SdrD) is a *Staphylococcus aureus* cell wall–anchored, calcium-binding adhesin member of the MSCRAMM Sdr subfamily that may contribute to bacterial adhesion and virulence. *S. aureus* is the most common cause of periprosthetic joint infection (PJI). Population-level distribution and sequence diversity of SdrD among clinical PJI isolates have not been systematically characterized, and the SdrD binding mechanism is still not well understood. To address these gaps, *sdrD* alleles were queried across 156 newly sequenced PJI isolates and compared to publicly available *S. aureus* genomes, and nucleotide– and protein-level phylogenies of the *sdrCDE* locus constructed. The SdrD crystal structure from *S. aureus* JH1 was determined, with solution small-angle X-ray scattering (SAXS) and molecular dynamics (MD) simulations, and assessment of conformational changes with calcium depletion. Three dominant *sdrD* subtypes were defined, associating with USA300, JH1, and TCH60; the JH1 *sdrD* subtype was predominant among PJI isolates. Structural studies showed that the conformation of individual domains and interdomain organization of the multidomain SdrD have limited flexibility in solution, and that the calcium-binding B domain retains its core fold under conditions of calcium depletion. Together, the findings presented support functional diversification among Sdr family members in mediating host attachment and inform a re-evaluation of the ligand-binding mechanism previously proposed for SdrD.

**AUTHOR SUMMARY:** *Staphylococcus aureus* is the leading cause of infections that develop around joint implants (periprosthetic joint infection, PJI). This bacterium has a large arsenal of surface proteins that allow it to stick to human tissues and implanted devices. This work focused on one such protein, SdrD, which has been linked to implant-associated infections but the structure and diversity of which among patients with PJI had not been well characterized. The genetic sequences of SdrD were analyzed across thousands of bacterial genomes, including those from patients with PJI. Distinct genetic variants of the protein were found, one of which was particularly common with PJI. The three-dimensional structure of SdrD was determined at atomic resolution and solution small-angle X-ray scattering (SAXS) and molecular dynamics used to study how it moves and responds to changes in its environment. Contrary to what was previously described, SdrD was shown to be relatively rigid. These findings change how SdrD’s mechanism of action should be considered, potentially informing design strategies to block bacterial attachment before infection takes hold.

## INTRODUCTION

*Staphylococcus aureus* ranks among the most clinically significant bacterial pathogens, with methicillin-resistant *S. aureus* (MRSA) responsible for an estimated 70,000 severe infections and 9,000 deaths per year In the United States (1). *S. aureus* is the causative organism for approximately one-third of orthopedic surgical site infections (2), and accounts for 25-33% of hip and knee periprosthetic joint infection (PJI) cases (3, 4). Despite advances in infection control, the burden of implant-associated *S. aureus* disease has proven difficult to reduce; PJI is associated with prolonged hospitalization, increased revision surgery rates and readmissions, and diminished 5-year survival (5, 6).

Central to *S. aureus* pathogenesis is its capacity for host tissue adhesion and surface colonization. Serine-aspartate repeat-containing protein D (SdrD) is an LPXTG-anchored adhesin displayed on the *S. aureus* surface (7–9). SdrD and related proteins—clumping factor A (ClfA), clumping factor B (ClfB), SdrC and SdrE—are grouped into a subfamily within the microbial surface components recognizing adhesive matrix molecule (MSCRAMM) family (10, 11). SdrD mediates adherence to keratinized epithelium and contributes to colonization and subsequent invasive infection (12, 13). By promoting adhesion and biofilm formation in protein-rich environments, SdrD, in concert with other adhesins, supports both initial colonization and transition to disease (14–16). SdrD has also been shown to enhance bacterial persistence in blood and reduce susceptibility to neutrophil-mediated killing (17). Desmoglein 1 (Dsg1), a desmosomal cadherin in the superficial epidermis, is a hypothesized target of SdrD (12, 18); however there has been no published attempt to co-crystallize SdrD bound Dsg-1. Investigation of the SdrD crystal structure and its associations with Dsg-1 would further inform this.

SdrD is prevalent across diverse *S. aureus* lineages, and SdrD-positive strains have been associated with invasive phenotypes, including bacteremia and musculoskeletal infections, including PJI (19–22). *In vivo* transcriptomic analysis of *S. aureus* PJI isolates revealed increased expression of *sdr* genes compared to the same strains grown in planktonic culture (23). This upregulation is likely not static as the *sdrD* locus in USA300 strain expands and contracts through recombination during *in vitro* growth and systemic infection, generating copy number heterogeneity in initially clonal populations (24). Among PJI specifically, within-patient genomic evolution involves adaptive mutations in adhesin-encoding loci (25), raising the possibility that SdrD variation contributes to persistence and treatment failure. Characterizing both the population-level distribution of *sdrD* in PJI and structural basis of SdrD function across lineages is therefore essential to further evaluating this adhesin.

Structures of several of *S. aureus* MSCRAMM subfamily members have been reported, such as collagen-binding protein (Cna) (26, 27), CflA (28, 29) and ClfB (30, 31); in addition, the related SdrG has been described from *Staphylococcus epidermidis* (32, 33). Structures of the N2-N3 domains of SdrG in complex with a fibrinogen-derived ligand peptide established a “dock, lock, and latch” (DLL) model of binding (32, 33). This model is supported by a structure of SdrE bound to a complement factor H peptide (34) and of the bone sialoprotein-binding protein (Bbp), and allelic variant of SdrE, in complex with a fibrinogen α peptide (35). The crystal structures of N2N3 and N2N3B1 domains SdrD from *S. aureus subsp. Aureus* TCH60 have also been reported (63). Despite these advances, key questions remain unresolved. Members of the Sdr subfamily share a common domain architecture primarily differing in B-domain number, yet it is unclear whether they universally employ a DLL-like binding mechanism through their conserved N2N3 region. If so, substrate specificity would be largely determined by the N2N3 binding groove, which has relatively low predicated sequence specificity; that the structural elements C-terminal to N3 required for the “lock” and “hatch” steps are the most divergent region among Sdr members, raises the possibility of alternative binding mechanisms. A further unresolved issue concerns the role of Ca^2+^-binding B domains; unlike the non-Ca^2+^-binding B domains of Cna (26), the B domains of SdrD (29), as well as the uro-adherence factor A (UafA) of *Staphylococcus saprophyticus* (36), suggest that its B domain participates directly in ligand recognition rather than functioning solely as a structural spacer (36). Whether SdrD’s B domains contribute similarly to ligand binding, and the mechanism that governs SdrD’s engagement with its proposed host ligand Dsg1 (12, 18), remain open questions.

Here, sequence diversity of SdrD was assessed across a collection of PJI isolates in comparison with publicly available *S. aureus* genomes, and findings used to inform selection of JH1 SdrD for characterization. Crystal structures of SdrD from *S. aureus* strain JH1, complemented by small-angle X-ray scattering (SAXS) in solution and molecular dynamics (MD) simulations, was determined, and conformational changes defined with Ca^2+^ depletion. Unexpected concordance was found between crystal structure and solution conformation of SdrD, including in interdomain organization, motivating re-evaluation of current models for SdrD–ligand recognition and binding.

## MATERIALS AND METHODS

### Collection of *S. aureus* isolates from PJI

*S. aureus* clinical isolates from prosthetic joint revisions collected at Mayo Clinic (Rochester, Minnesota) between 1999 and 2022 were studied. Isolate sources include either synovial fluid collected prior to surgery or intraoperatively, tissue collected intraoperatively, or sonicate fluid collected from resected orthopedic hardware. Synovial fluid was cultured using BACTEC bottles (BD Diagnostic Systems) incubated on a BACTEC FX 9240 instrument (BD Diagnostic Systems) or on standard aerobic and anaerobic media. Tissues were homogenized using a Seward Stomacher 80 Biomaster (Seward Inc., Port St. Lucie, FL) in 5 mL of brain heart infusion broth for 1 minute. Between 1999 and 2015, homogenate was inoculated onto standard aerobic and anaerobic media and incubated at 35-37 °C in 5-7% CO_2_ for 5 days or anaerobically for 14 days, respectively. Between 2016 and 2022, homogenate was inoculated into BACTEC Plus Aerobic/F and BACTEC Lytic/10 Anaerobic/F bottles and bottles incubated in a BACTEC FX9240 instrument for 14 days or until positive. For sonication cultures, prosthesis components were placed in solid containers in the operating room. In the laboratory, 400 mL sterile Ringer’s solution was added to each container, and containers subjected to vortexing for 30 seconds and sonication for 5 minutes, followed by another round of vortexing (37). Between 2001 and 2005, 0.5 ml sonicate fluid was plated onto aerobic and anaerobic sheep blood agar plates, incubated at 35-37°C in 5-7% CO_2_ for 5 days or anaerobically for 7 days, respectively. Between 2006 and 2022, a 100-fold concentration by centrifugation was performed, and 0.1 ml of concentrated sonicate fluid plated on aerobic and anaerobic solid media, with 2-to-4-day incubation and 14-day incubation, respectively. *S. aureus* isolates were identified using standard microbiology techniques, and frozen at –80 °C in a Microbank^®^ cryovial (Pro-Lab Diagnostics) until the time of sequencing.

### Sequencing of PJI *S. aureus* genomes

Isolates were streaked from frozen onto TSB Sheep Blood Agar plates and grown at 37°C for 18 hours (h). Cultures were suspended in 500 μL of TE buffer to the density of a 4 McFarland standard. Suspended cells were incubated with 100 μL 0.5 mg/mL lysostaphin for 3 h at 37°C. To remove RNA, 15 μL of RNAse (Invitrogen Purelink 20 mg/mL 12091021) was added and incubated for 15 minutes (min). DNA was extracted using a Promega Maxwell® RSC and Promega Maxwell® Tissue DNA Kit (Promega, AS1030). To remove remaining beads, extracts were cleaned using a Monarch® Spin PCR & DNA Cleanup Kit (5 μg) (NEB #T1130). DNA concentrations were read using a Qubit and QuantiFluor® dsDNA System Kit (E2671). Genomic DNA was sequenced by SeqCoast Genomics on an Illumina platform. Libraries were prepared and sequenced to generate paired-end short reads, yielding a target output of approximately 400 Mbp (2.7 million reads) per isolate, a sequencing depth corresponding to an estimated mean coverage of 130× per genome, based on the expected *S. aureus* genome size of ∼2.8–3.0 Mbp. Eight genomes previously sequenced by Masters et al. were also included (PRJNA700676) (38).

### *S. aureus* PJI genome alignment

All alignments and downstream phylogenetic analyses were performed using an aarch64-apple-darwin20 platform (Apple Silicon, macOS). Code is available on GitHub. Raw paired-end reads were quality trimmed using Trimmomatic (v0.39) in paired-end mode (PE). Reads were processed with a sliding window of 4 bases requiring a minimum average Phred quality score of 20 and reads shorter than 50 bp post-trimming discarded. Only read pairs in which both reads survived trimming (paired output) were retained for genome assembly; unpaired surviving reads were discarded. Trimmed paired-end reads were assembled using SPAdes (v3.15). Chromosomal genome assembly was performed using the –-isolate flag, which is optimized for high-coverage single-isolate sequencing data. To recover plasmid sequences, a separate assembly was performed on each sample using plasmidSPAdes (--plasmid), which is designed to detect and assemble plasmid contigs from Illumina short-read data.

### Publicly available *S. aureus* sequences

Dehydrated closed *S. aureus* contigs were downloaded from NCBI using a command line and rehydrated; on September 23^rd^, 2024, this amounted to 6,607 contigs (genomes and plasmids). There were 2,202 unique BioSample IDs, suggesting that some genomes were duplicates, perhaps due to the presence of both draft and finalized genome assemblies. As SAMD00180473 was more genetically like *S. argenteus* or *S. schweitzeri* than *S. aureus*, it was excluded from analysis.

### Core genome multilocus sequence typing and phylogenetic analysis

Core genome multilocus sequence typing (cgMLST) was performed using chewBBACA (v3.x) with the scheme (Scheme 20) comprising 1,716 alleles. Prior to allele calling, the scheme was prepared using the *S. aureus* Prodigal training file obtained from the chewBBACA GitHub repository. Individual genome assemblies in fasta format were organized into a single directory, and allele calling performed across all genomes using the AlleleCall module with 8 CPU threads. Following allele calling, loci were filtered using the ExtractCgMLST module, retaining only those present in ≥95% of genomes (to minimize the impact of missing data on downstream analyses). Genomes with greater than 10% missing loci were subsequently identified and removed using R Studio (version 4.3.2). Multilocus sequence types (STs) were assigned using MLST with the *S. aureus* (*saureus*) scheme. A minimum spanning tree was constructed from the filtered cgMLST profiles using GrapeTree with the MSTreeV2 algorithm. Trees were visualized and annotated with MLST data in R using the ggtree package.

### *sdr* sequence queries using Gene Allele Mutation Microbial Assessment

Gene Allele Mutation Microbial Assessment (GAMMA) (39) was used to query for gene sequences against multifasta files containing all publicly available *S. aureus* contigs and clinical PJI isolates. DNA sequence files for *sdrC*, *sdrD*, and *sdrE* were used from *S. aureus* strain USA300, from NCBI (Assembly: <u>ASM15366v1</u>). The –f tag was included to output the fasta sequences for query results.

### Sequence alignment and phylogeny assessment

The fasta file of queried *sdr* outputted from GAMMA was inputted into R Studio to create alignments and phylogenetic trees. The DECIPHER (version 2.30.0) R package was used to create alignments of *sdr* (40, 41). After converting the alignment into a phyDat matrix, the distance matrix was calculated using the dist.ml command from the phangorn package (42, 43). A Newick tree file was created from the distance matrix using the nj command from ggtree. An unrooted tree was then created using the ggtree “unrooted” setting, which uses the “daylight” method as a default. Clades and tips were labeled using ggtree functionalities.

### Cloning, protein purification and crystallization

Gene fragments encoding N2N3 (residues S244–S562), B1 (residues G564–Y687), N2N3B1 (residues S244–Y687), N2N3B1B2 (residues S244–Y798), and N2N3B1B2B3 (residues S244–Y908) were PCR-amplified and cloned into a pMCSG53 vector. The N574A point mutation in the N2N3B1 construct was generated by site-directed mutagenesis and cloned into the same vector. Cloning into pMCSG53 introduced an N-terminal His₆-tag followed by a TEV protease cleavage site.

For protein expression, plasmids were transformed into *Escherichia coli* BL21 (DE3)-Gold cells (Stratagene) using heat shock. Transformed cells were grown overnight at 37°C in 100 mL LB medium containing 150 µg/mL ampicillin. 10 mL of overnight culture was inoculated into 1 L LB medium supplemented with 150 µg/mL ampicillin and grown at 37°C. At OD₆₀₀ 0.8–1.0, protein expression was induced with 0.4 mM IPTG, followed by incubation at 18°C for 16 h. Cells were harvested at 4°C at 5,000 × g for 20 min.

For construct purification, cell pellets were resuspended in lysis buffer (50 mM HEPES pH 8.0, 500 mM NaCl, 20 mM imidazole, 10 mM β-mercaptoethanol [BME], and 5% [v/v] glycerol), lysed by sonication, and clarified by centrifugation. Proteins were purified by immobilized metal affinity chromatography (IMAC) using 5 mL Ni²⁺-Sepharose (Cytiva) packed in a Flex-Column operated on a Vac-Man vacuum system (Promega). After equilibration with lysis buffer, clarified lysate was loaded onto the resin, washed with 20 column volumes of lysis buffer, and eluted using buffer containing 50 mM HEPES pH 8.0, 500 mM NaCl, 500 mM imidazole, 10 mM BME, and 5% glycerol. Eluted proteins were incubated with His-tagged TEV protease (1:20 molar ratio of TEV to target protein) and dialyzed overnight at 4°C against lysis buffer. To remove the cleaved His-tag and His-tagged TEV protease, s a second IMAC purification step using Ni²⁺-Sepharose was performed under the same buffer conditions, and the flow-through containing the tag-free target proteins collected. Proteins were then concentrated using 10 kDa molecular-weight-cutoff centrifugal concentrators (Merck-Millipore), and subsequently buffer-exchanged into protein buffer (20 mM HEPES pH 7.5, 150 mM NaCl, 1 mM TCEP). Finally, proteins were analyzed by 4–20% SDS–PAGE, concentrated to 30 mg/mL, and stored at −80°C.

Purified proteins (30 mg/mL) were screened for crystallization with a Mosquito nanoliter liquid handler (TTP LabTech) using the sitting drop vapor diffusion methods. Crystallization screening was performed using MCSG1-4 (Anatrace) screens at 16°C. Each crystallization droplet containing 0.4 µL of each protein and precipitating agent was equilibrated against a reservoir containing 140 µL of precipitating solution. Three proteins, N2N3, N2N3B1 and N2N3B1B2, were crystallized under several conditions. Representative crystallization conditions included 0.2 M calcium chloride and 20% (w/v) PEG 3350 for N2N3; 0.16 M magnesium chloride, 0.08 M Tris–HCl, pH 8.5, and 24% (w/v) PEG 4000 for N2N3B1; and 0.1 M Tris–HCl, pH 7.0, and 20% (w/v) PEG 2000 MME for N2N3B1B2. Crystals were harvested, treated with cryoprotectant solutions (25% [v/v] glycerol or ethylene glycol in the corresponding mother liquor), and flash-frozen in liquid nitrogen before X-ray diffraction data collection.

### X-ray diffraction and structural determination

X-ray diffraction data were collected at 100 K from cryocooled crystals at the 17-ID-2 or 19-ID beamlines of National Synchrotron Light Source II at Brookhaven National Laboratory. Intensities were integrated, scaled, and merged with the HKL-3000 program suite (44) or Xia2 (45). The N2N3B1 structure was first resolved by using molecular replacement (MR) method (46), with the homolog SdrD structure (PDB ID: 4JDZ) as a search template. The individual domain (or domains) from the first structure was (or were) used as a template (or templates) in the following structural determination of N2N3 and N2N3B1B2 by using MR. The final models were completed following iterative rounds of model rebuilding using Coot (27), with restrained refinement using Phenix.refine (47), and were validated with using Molprobity (48) (Table 1).

**TABLE 1.** Crystallographic data.

| Data collection statistics | N2N3 | N2N3B1 | N2N3B1B2 |
| --- | --- | --- | --- |
| Space group | *I*222 | *P*2<sub>1</sub>2<sub>1</sub>2<sub>1</sub> | *P*2<sub>1</sub> |
| Unit cell (Å, °) | *a* = 70.52<br>*b* = 73.15<br>*c* = 198.7 | *a* = 43.05<br>*b* = 72.83<br>*c* = 148.5 | *a* = 59.10<br>*b* = 58.40<br>*c* = 94.30<br>*b* = 101.3 |
| MW Da (number of amino | 35,181 (319)<sup>1</sup> | 48,673 (444)<sup>1</sup> | 60,716 (555)<sup>1</sup> |
| Mol (AU) | 1 | 1 | 1 |
| Wavelength (Å) | 0.979 | 0.979 | 0.979 |
| Resolution (Å) | 1.92-37.0 | 1.80-30.0 | 2.00-27.2 |
| Number of unique | 39,344 (1,851)<sup>2</sup> | 44,004 (11,542)<sup>3</sup> | 42,903 (2,157)<sup>4</sup> |
| R<sub>merg</sub> (%) | 5.8 (84.6)<sup>2</sup> | 8.5 (95.2)<sup>3</sup> | 12.0 (100.6)<sup>4</sup> |
| R<sub>pim</sub> (%) | 2.9 (50.1)<sup>2</sup> | 7.2 (86.5)<sup>3</sup> | 7.6(66.1)<sup>4</sup> |
| CC ½ | 0.997 (0.549)<sup>2</sup> | 0.994 (0.403)<sup>3</sup> | 0.995 (0.645)<sup>4</sup> |
| Completeness (%) | 98.8 (94.3)<sup>2</sup> | 98.0 (96.2)<sup>3</sup> | 99.6 (99.8)<sup>4</sup> |
| Redundancy | 5.0 (3.4)<sup>2</sup> | 13.4 (2.9)<sup>3</sup> | 3.4 (3.2)<sup>4</sup> |
| I/(σ) | 29.2 (1.2)<sup>2</sup> | 9.40 (0.89)<sup>3</sup> | 8.4 (1.2)<sup>4</sup> |
| Solvent content (%) | 66.6 | 48.5 | 53.5 |
| **Phasing** |  |  |  |
| Resolution range (Å) | 37.0 – 3.00 | 30.0 – 3.00 | 27.5 – 2.00 |
| Correlation coefficient (%) | 49.94<sup>3</sup> | 43.55<sup>3</sup> | 64.00<sup>3</sup> |
| **Refinement** |  |  |  |
| Resolution range (Å) | 37.0 – 1.92 | 30.0 – 1.80 | 27.5 – 2.00 |
| Number of reflections | 39,302 | 43,927 | 42,858 |
| Completeness (%) | 99.3 | 99.8 | 99.5 |
| R<sub>work</sub>/R<sub>free</sub> (%) | 21.4/24.5 | 18.0/21.1 | 21.0/23.5 |
| No. of Atoms | 2,574 | 3,759 | 4,604 |
| Bond lengths (Å) | 0.007 | 0.004 | 0.002 |
| Bond angles (deg) | 0.837 | 0.656 | 0.483 |
| B-factors (Å<sup>2</sup>) (main/side) | 54.4/57.1 | 33.0/36.9 | 34.2/37.7 |
| Wilson B-factor (Å<sup>2</sup>) | 36.9 | 26.4 | 29.6 |
| **Molprobtity validation** |  |  |  |
| Ramachandran outliers (%) | 0.00 | 0.00 | 0.00 |
| Ramachandran Favored (%) | 96.83 | 97.43 | 96.75 |
| Rotamer outliers (%) | 0.35 | 1.55 | 0.41 |
| Clashscore | 3.49 | 2.36 | 2.11 |
| **PDB ID** | 10PA | 8VDK | 10PS |

<sup>1</sup>Not including N-terminal three vector derived residues, SNA<sup>2</sup>Last resolution bin, 1.95-1.92 Å<sup>3</sup>Last resolution bin, 1.85-1.80 Å<sup>4</sup>Last resolution bin, 2.03-2.00 Å<sup>5</sup>Molecular replacement method

### Solution biological small-angle X-ray scattering

Solution biological small-angle X-ray scattering (SAXS) was performed at beamline 12-ID-B of the Advanced Photon Source (APS) at the Argonne National Laboratory. The wavelength of X-ray radiation was set to 0.932 Å. Scattered X-ray intensities were measured using an Eiger2 9M detector (DECTRIS LLC). The sample-to-detector distance was set such that the detecting range of scattering momentum transfer *q* was 0.004–0.8 Å^-1^. The scattering momentum transfer q is defined as q=4πsin(θ)/λ, where λ represents the X-ray wavelength and 2θ corresponds to the Bragg angle. To remove effects of potential impurities, samples were measured using an inline AKTA micro FPLC setup with a Superdex 200 Increase 5/150 GL size exclusion column (SEC) (Figure S2). Samples were passed through the FPLC column and fed to a flow cell, which consists of a cylindrical quartz capillary 1.5 mm in diameter with a 10 µm wall thickness, for SAXS measurement. The flow rate was set at 0.2 mL/min; SAXS images were collected every second with an exposure time of 0.3 seconds. 2-D SEC-SAXS images were converted to 1-D SAXS (I(q) vs q) curves through azimuthally averaging after solid angle correction and then normalization with the intensity of the transmitted X-ray beam flux, using the beamline software package matSAXS (https://12idb.xray.aps.anl.gov/Software_Processing.html). To further remove the influence of possible impurities, 1D SEC-SAXS data were processed using evolving factor analysis in software BioXTAS RAW (version 2.3.1) (49). Evolving Factor Analysis (EFA) is widely used in SEC-SAXS to determine how many distinct scattering species are present as a function of elution time, to identify where each species appears and disappears in the chromatogram, and to deconvolute and obtain pure SAXS spectra and concentration profiles.

SAXS profiles were subjected to further quantitative analyses. The radius of gyration (*R*g), a size parameter, was calculated on a low *q* region using the Guinier equation, i.e. 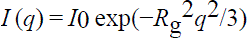, where *I*0 is the forward scattering. Guinier plots, ln[I(q)] *versus* q^2^, were additionally used to assess sample uniformity and data quality. The linear Guinier regions observed for the SEC-SAXS datasets of the *S. aureus* SdrD proteins (Figure S3) indicated excellent size monodispersity and high data quality, supporting reliable real space analyses. For samples whose atomic structure are available, obtained either from crystallography or computational modeling, the experimental SAXS dataset was fitted with the atomic structure using program CRYSOL (50) from the ATSAS package (51). CRYSOL calculates the SAXS profile of the atomic structure and adds proper amounts of an hydration layer to achieve the best fit (Figure S4, for example). The goodness-of-fit was judged by the value of Chi2 (χ^2^), defined as follows, 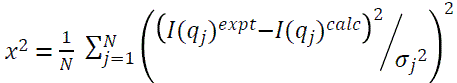, where N is the total number of data points, and σj is the experimental error on the SAXS intensity at the jth data point. Crystallographic atomic structures using in the CRYSOL SAXS fittings were partially (for constructs B1, N2N3, and N2N3B1) or fully (for N2N3B1B2) adopted from the crystal structure of N2N3B1B2 solved in this work (pdb ID: 10PS) depending on whether all residues were resolved or some were missing in the crystal structures.

The pair distance distribution function (PDDF, *P*(*r*)) that is the inverse Fourier transform of X-ray scattering data and represents a weighted histogram of atomic-pair distances in real space. The maximum molecular dimension (Dmax) can be estimated from PDDFs as the distance at which P(r) approaches zero, i.e., P(r>=Dmax) = 0. Empirically, Dmax values derived from PDDFs may carry uncertainties of 10% or more, with larger errors typically observed for elongated particles or asymmetric PDDFs exhibiting extended tails. The program GNOM (52) from the ATSAS package was employed to calculated PDDF profiles from SAXS profiles, with the program SolX3 (53) used to compute PDDF profiles directly from atomic structures, if available.

Three-dimensional real space molecular envelopes were reconstructed from SAXS data using DAMMIN/DAMMIF (54) and GASBOR (55) from the ATSAS package, and program DENSS (56). These methods produced consistent SAXS molecular envelopes; therefore, only DAMMIN/DAMMIF results are reported here. DAMMIF is a fast version of DAMMIN, with DAMMIN providing more user options. Both programs employ simulated annealing algorithm to search structures that best fit SAXS data from an ensemble of densely packed beads (or dummy atoms).

For each SAXS dataset, ten to 15 DAMMIF independent reconstructions were first performed using SAXS data up to *q* of ∼0.35 Å^-1^. Resulting models were aligned and averaged, followed by a DAMMIN refinement on a confined search space defined by the DAMMIF ensemble. This combined DAMMIF/DAMMIN workflow yields an averaged DAMMIF model (also referred to a consensus model based on bead occupancy over all DAMMIF reconstructions), a most-probable DAMMIF model (with the least RMSD relative to the rest reconstructions), and a DAMMIN-refined model (51). Both the most-probable and refined models fit SAXS data well; however, refined models typically use larger bead sizes, therefore with lower effective resolution. Accordingly, results with the most-probable models are preferentially reported here, except where the averaged RMSD among the reconstructions exceeded 30 Å, in which case results with the refined model are reported.

### Thermal shift assays

Thermal shift assays were conducted using a Bio-Rad CFX Connect™ Real-Time PCR Detection System (Bio-Rad Laboratories). Purified SdrD constructs comprising the N2N3, B1, N2N3B1, N2N3B1B2, and N2N3B1B2B3 domains, and the N2N3B1_N57A mutant, were assayed at a final concentration of 40 µM in buffer containing 20 mM HEPES (pH 7.5), 150 mM NaCl, and 1 mM TCEP. Proteins were incubated in the presence or absence of peptide (1 mM) and subsequently mixed with 5X Sypro Orange fluorescent dye. Thermal denaturation was monitored by increasing the temperature from 25 to 95°C in 1°C increments, with fluorescence data collected as two endpoint readings per 1 min cycle. A predefined method implemented in the CFX software was used to calculate the first derivative of the fluorescence-based denaturation curve. The apparent melting temperature (Tm) was defined as the temperature corresponding to the maximum rate of fluorescence change, determined from the local minimum of the first derivative curve [−d(fluorescence)/dT], which reflects the midpoint of protein unfolding.

### Molecular Dynamics simulation

Molecular Dynamics (MD) simulations were conducted on the B1 domain to examine the difference in stability and secondary structure composition with and without Ca^2+^; Equilibrium and Replica Exchange with Solute Tempering (REST2) was used (57). MD simulations were performed using the CHARMM36m forcefield (58). NAMD was used carry out the MD simulations (59). VMD (60) was used to solvate the B1 domain in TIP3P water with 2 nm padding in each direction (61). Equilibrium MD simulations were run for 300 nanoseconds (ns) at 300 and 450 K in the NPT ensemble, following an initial equilibration period in the NVT ensemble. The NPT ensemble was maintained using Langevin dynamics and a Nose-Hoover barostat, with pressure set to 1 atm and a decay time of 100 fs. The timestep was set to 2 femtoseconds (fs), and the particle-mesh Ewald method used to calculate full electrostatic interactions. REST2 simulations were performed for an additional 500 ns, using the equilibrium simulation as the initial coordinates. Bonds between heavy atoms and hydrogens were constrained using the SHAKE algorithm (62). Periodic boundary conditions were used for all simulations. The REST2 simulation used 16 replicas from an effective temperature of 300 to 450 K to enhance sampling of the conformational landscape. Exchanges were attempted every 1 picosecond; a scaling factor of 0.7 was used for protein water interactions. Theoretical SAXS for the conformational ensembles of the B1 domain with and without Ca^2+^ were calculated using SolX3 (53).

## RESULTS

### Phylogeny of *sdrD* from *S. aureus* references reveal three *sdrD* sequence subtypes

SdrD is enriched among bone and implant-associated *S. aureus* strains (19–22), yet whether this reflects clonal expansion of specific linages or convergent retention of the locus across diverse genetic backgrounds is unknown. This work considered *sdrD* genes from all publicly available closed *S. aureus* contigs; at the time of analysis, this amounted to 6,607 publicly available closed contigs, 4,152 of which were unique *S. aureus* genomes and plasmids from 2,202 unique *S. aureus* BioSamples. Using GAMMA, a BLAT-based genomic search tool, *sdrD* was queried against the library of publicly available isolates using the *sdrD* sequence from USA300, a well-characterized and frequently used *S. aureus* laboratory and genomic reference strain. USA300 *sdrD* returned 1,300 sequences, most 4,146 nucleotides in length. It was suspected that some instances of *sdrD* might have been missed, so the search was repeated with JH1, a secondary reference strain to search for potentially additional hits. JH1 *sdrD* returned 1,442 sequences, most 4,014 nucleotides in length. These fundings suggested that *sdrD* might have multiple nucleotide subtypes, and that they diverged from one another to the extent that GAMMA’s blat-based search could not identify a different subtype than what it was querying for. To further explore this, a collection of well annotated reference strains was used to build a phylogeny of *sdrD* sequences. In addition to USA300 and JH1, COL, Newman, NCTC8325, MW2, JH9 and N315, as well as TCH60 which was previously used to characterize the crystal structure of SdrD (63), were included. Using the USA300 and JH1 sequences for *sdrD*, *sdrD* was queried against references, and the queried returns used to assemble a phylogenetic tree. USA300, COL, Newman, NCTC8325, and MW2 formed one node, separate from JH1, JH9 and N315, which formed a second node, which are referred to as *sdrD_USA300* and *sdr_JH1 nodes*, respectively. TCH60 was most divergent from other sequences and formed a node on its own (‘*sdrD_TCH60* node’), distinct for the other reference strains (**Figure 1A).**

**Figure 1.**
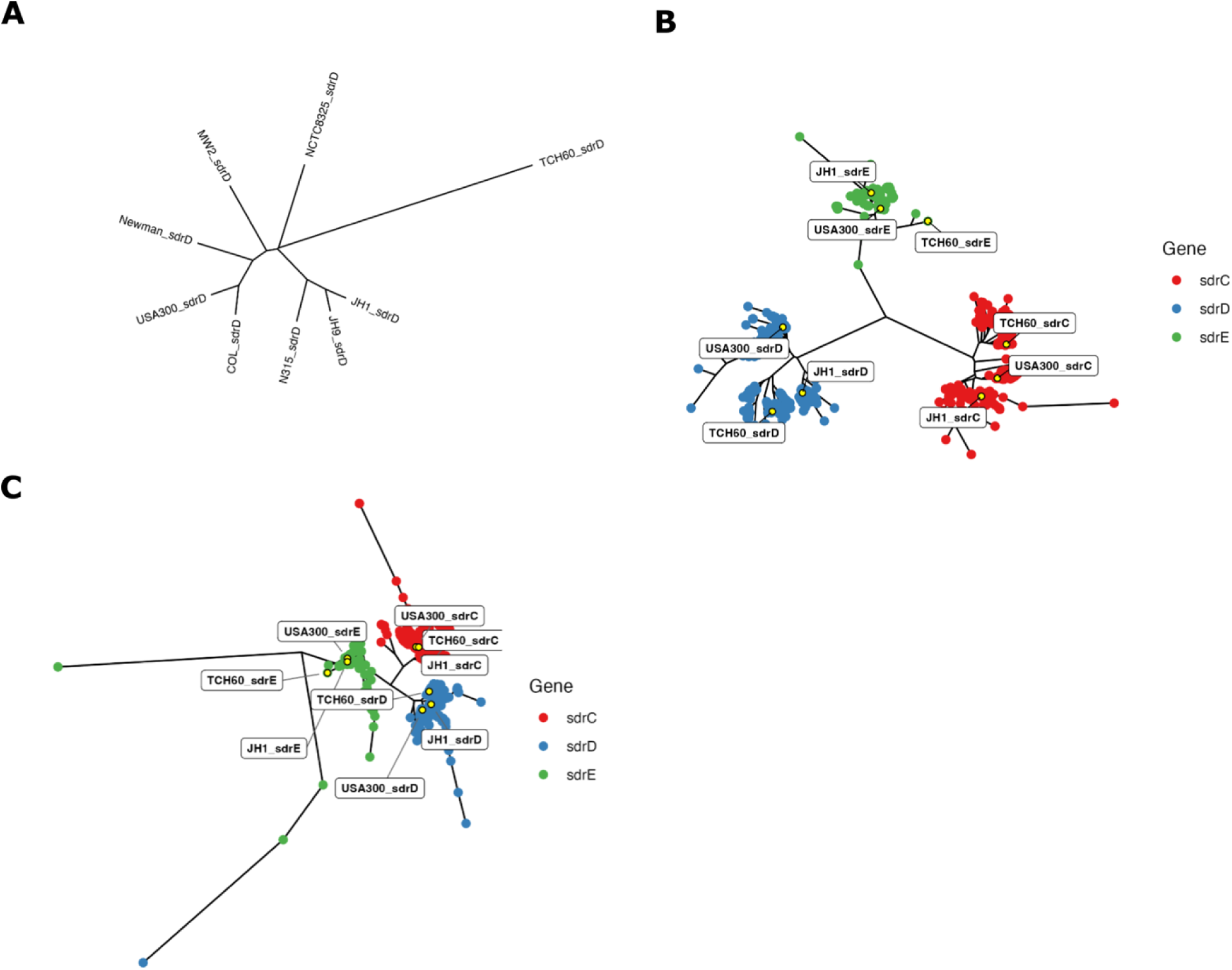
Phylogenetic analysis of SdrCDE nucleotide and amino acid sequences from reference strains and a large public *S. aureus* library reveals three distinct SdrD sequence subtypes. (A) Unrooted neighbor-joining phylogenetic tree of *sdrD* nucleotide sequences from nine *S. aureus* reference strains (USA300, COL, Newman, NCTC8325, MW2, JH1, JH9, N315, and TCH60). Three distinct nodes are observed: a USA300-associated node (comprising USA300, COL, Newman, NCTC8325, and MW2), a JH1-associated node (comprising JH1, JH9, and N315), and a TCH60 node. (B) Unrooted neighbor-joining phylogenetic tree of *sdrC* (red), *sdrD* (blue), and *sdrE* (green) nucleotide sequences derived from queries against a large public *S. aureus* library (*sdrC*: n=3,350; *sdrD* USA300 query: n=1,300; *sdrD* JH1 query: n=1,442; *sdrE*: n=2,791), along with sequences from reference strains. Sequences exceeding 5,000 nucleotides were excluded. USA300, JH1 and TCH60 associate with distinct nodes and recapitulate the three-node structure observed in (A). (C) Unrooted neighbor-joining phylogenetic tree of translated SdrC (red), SdrD (blue), and SdrE (green) amino acid sequences from the public *S. aureus* library. Three SdrD protein nodes are observed, associated with USA300, JH1, and TCH60 reference sequences, consistent with the nucleotide-level phylogeny. One SdrD sequence clustered with an SdrE offshoot.

### Phylogeny of *sdrCDE* from large *S. aureus* library

Based on the observation that the reference strains were associated with three *sdrD* nodes, the phylogenetic distribution of *sdrD* was investigated using a larger collection of isolate sequences. *sdrD* sequences from JH1 and USA300-based queries from the large public *S. aureus* library were included, as were *sdrC* and *sdrE* sequences, since *sdrCDE* genes are thought to have arisen from gene duplication events, to add context to the sequence divergence observed in **Figure 1A**. In addition to the JH1 and USA300-based *sdrD* queries described above, gene sequences for *sdrC*, *sdrD*, and *sdrE* from USA300 were queried against the public *S. aureus* library. USA300 sequences for *sdrC* and *sdrE* returned 3350 and 2791 sequences, respectively. Given that *sdrD* was the primary interest, it was decided that the USA300 query had returned a sufficiently high number of *sdrC* and *sdrE* hits that iterative query with more isolate sequences was not needed. Sequences returned by the queries were aligned along with *sdrD* sequences from USA300, JH1, COL, Newman, NCTC8325, MW2, JH9 and N315, and *sdrC* and *sdrE* sequences from USA300 and JH1 and used to create an unrooted neighbor joining tree (**Figure 1B**). Sequences longer than 5,000 nucleotides were excluded from downstream alignment and tree creation. *sdrC* sequences are plotted in red, *sdrD* sequences in blue and *sdrE* sequences in green. The three genes each formed distinct nodes. As with the smaller phylogenetic analysis (**Figure 1A**), *sdrD* formed three primary nodes, associated with USA300, JH1, and TCH60. *sdrC* similarly formed three nodes associated with TCH60, USA300 and JH1. *sdrE* formed a single primary node which included USA300 and JH1, with TCH60 an offshoot.

To assess divergence between protein sequences, nucleotide sequences returned from the *S. aureus* public library were translated to amino acid sequences using DECIPHER. Protein sequences were plotted as an unrooted neighbor joining tree, with SdrC, SdrD, and SdrE sequences in red, blue, and green, respectively (**Figure 1C**). As observed with nucleotide sequences, each protein formed a distinct node, except for one SdrD sequence, which associated with offshoot SdrE proteins. SdrD protein sequences formed three primary nodes, each associated with JH1, USA300 or TCH60. This suggests that SdrD TCH60 represents only a subset of *S. aureus* isolates – to understand the function of SdrD in *S. aureus*, the structures of alternative SdrD need to be defined.

### Phylogeny of *sdrCDE* from PJI isolates

While the *S. aureus* public library represents a large breadth of *S. aureus* isolates, it does not include many PJI isolates. Accordingly, the above analysis was repeated with a collection of 156 *S. aureus* isolates from PJI, each collected from a unique patient. **Figure 2A** shows a phylogenetic tree for the isolates’ genomes analyzed using cgMLST, with the number of alleles differing between each isolate shown in **Figure 2B**. *sdrCDE* sequences from USA300, JH1, and TCH60 were queried against each PJI genome. USA300, JH1 and TCH60 sequences were chosen based on the results shown in **Figure 1B** and **1C**. *sdrC* was found in 153/156 isolate sequences using the USA300 *sdrC* sequence as a query, but only 136 and 144 when using JH1 or TCH60 s*drC* query sequences, respectively. *sdrD* was found in only 44/156 isolate sequences when using the USA300 s*drD* sequence as a query; query with JH1 and TCH60 *sdrD* returned 79 and 53 sequences, respectively. Querying with USA300 *sdrE* returned 104/156 isolate sequences, with JH1 and TCH60 returning 112 and 53, respectively. Queried sequences for *sdrC*, *sdrD*, and *sdrE* from 156 PJI isolates along with the references used above were used to create an unrooted neighbor joining tree with *sdrC*, *sdrD* and *sdrE* sequences are plotted in red, blue and green, respectively (**Figure 2C**). In summary, BLAT-based searches and phylogenetic clustering show that JH1 is best representative of SdrD for PJI isolates. Given the importance of SdrD for virulence and its previously documented upregulation during PJI, subsequent analysis focused on structural characterization of JH1 SdrD.

**Figure 2.**
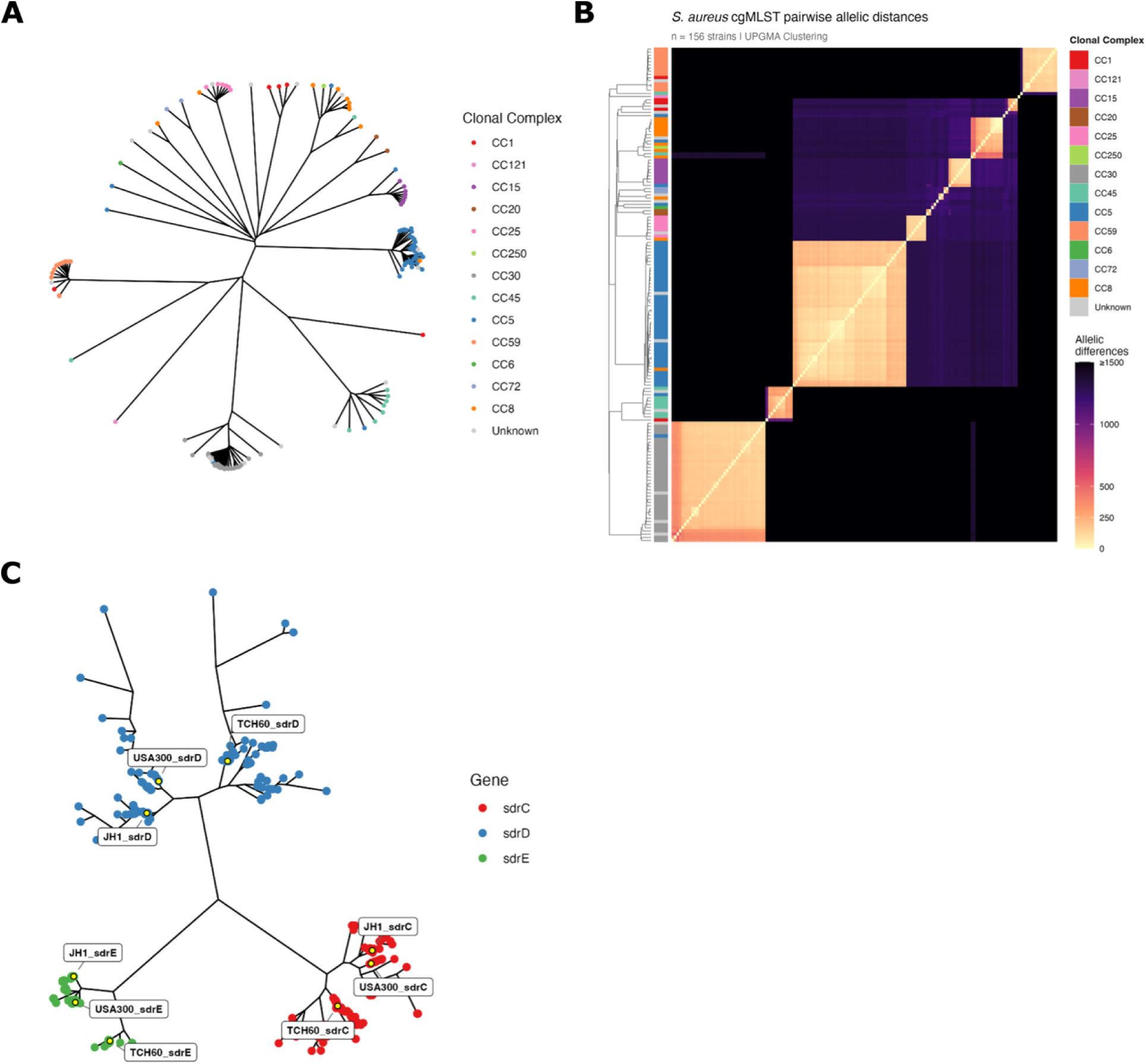
Phylogenetic analysis using core genome multilocus sequence typing (cgMLST) and *sdrCDE* sequences from *Staphylococcus aureus* isolates from periprosthetic joint infection (PJI). (A) Unrooted neighbor-joining phylogenetic tree of 156 *S. aureus* isolates recovered from PJI, each from unique patients, constructed using core genome multilocus sequence typing (cgMLST). (B) Pairwise cgMLST allelic distance matrix illustrating the number of differing loci between each pair of PJI isolates. (C) Unrooted neighbor-joining phylogenetic tree of *sdrC* (red), *sdrD* (blue), and *sdrE* (green) nucleotide sequences in the 156 PJI isolates. using USA300, JH1, and TCH60 as query sequences. *sdrC* was detected in 153 isolates (USA300 query), *sdrD* in 44 (USA300), 79/156 (JH1), and 53/156 (TCH60), and *sdrE* in 104 (USA300), 112 (JH1), and 53 (TCH60). The JH1 *sdrD* query returned the greatest number of *sdrD* sequences among PJI isolates, supporting JH1 as the most representative reference for SdrD structure-function studies in the context of PJI.

### N2N3 domains – relative domain-domain movement and Ca^2+^-binding sites

SdrD comprises of multiple domains including N-terminal signal peptide, an “A” region (N2–N3 subdomains) that binds host ligands, B-repeats, a long SD-repeat stalk, and a C-terminal LPXTG motif for Sortase A-mediated cell wall anchoring (**Figure 3A**) (9). N2 and N3 domains are the two stably folded domains in the in A-region of SdrD (**Figure 3A**). In multiple constructs expressed, the three crystal structures determined contain N2N3, including N2N3 itself, N2N3B1 and N2N3B1B2 (**Figure 3C-E**). Analysis and comparison of these structures reveals N-domain structural details and dynamic features.

**Figure 3.**
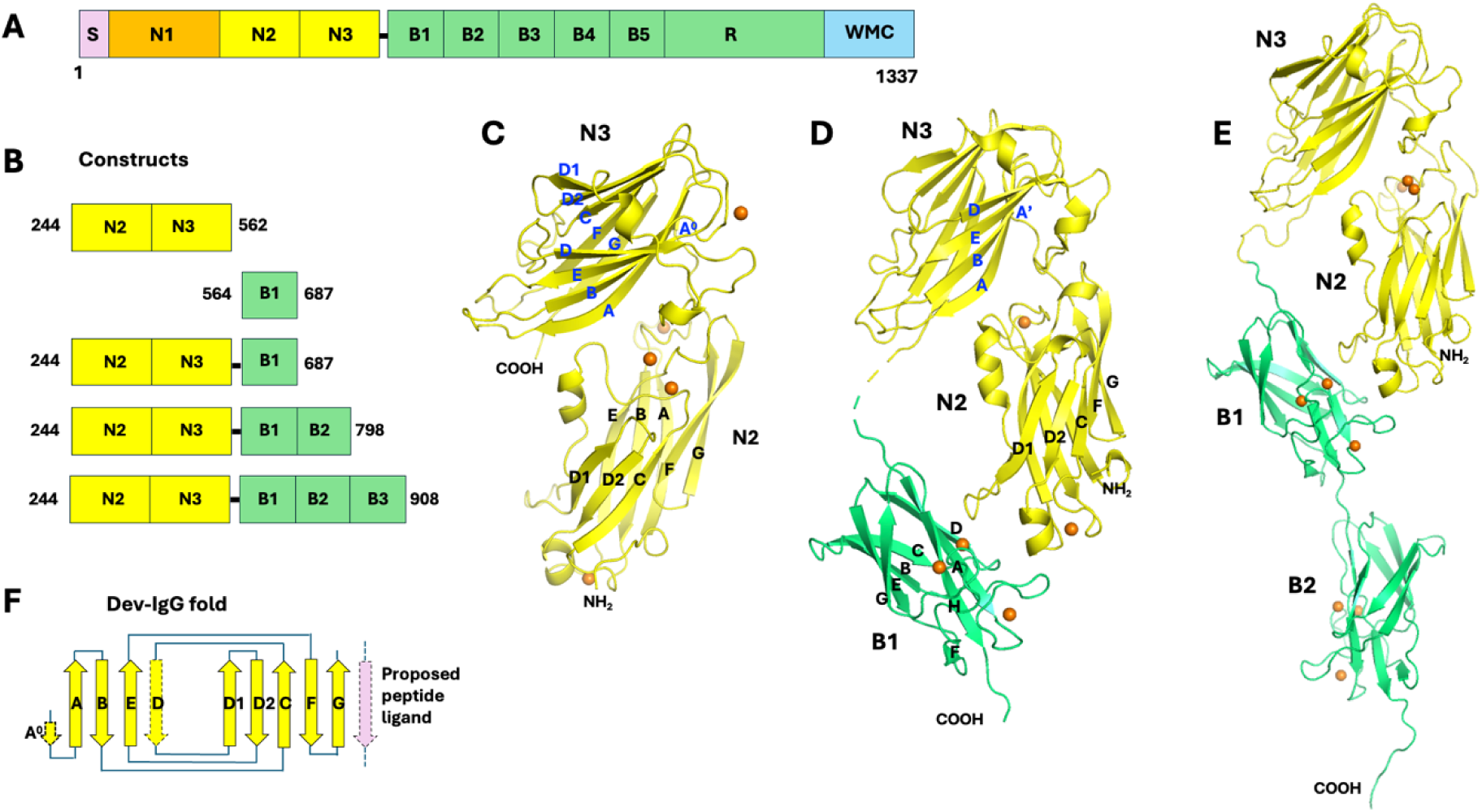
*Staphylococcus aureus* SdrD molecular structure. (A) *S. aureus* SdrD domain organization. The molecule includes a signal peptide (S), a so-called “A-region” including a predicted unstructured N1 “domain” and structured N2 and N3 domains, five B repeats (domains), a SD-repeat region (R), and a C-terminal anchoring module, including a wall/sorting signal (LPXTG motif) (W), transmembrane segment (M) and cytoplasmic tail (C). (B) The *S. aureus* SdrD constructs used in this work. (C) A ribbon diagram of crystal structure N2 and N3 domains (N2N3). All ß-strands are labeled. Ca^2+^ ions are displayed as small orange spheres. (D) A ribbon diagram of crystal structure of N2N3B1 domains, N2 and N3 is shown in yellow and B1 in green. All ß-strands of B1 domain are labeled. Strands on one ß-sheet of N2 and one ß-sheet of N3 domains are labeled only for their orientation. (E) A ribbon diagram of crystal structure of N2N3B1B2 domains. The color scheme for N and B domains is the same as in Figure 1D. Structures in all ribbon diagrams are not on the same scale. (F) The topology scheme of Dev-IgG fold found in N2 and N3 domains. The possible peptide ligand binding on N3 domain is illustrated by a pink strand antiparallel with edge G strand. A peptide segment after N3 domain may form a latch in the position of D strand of N2 according to “dock, lock and latch (DLL)” substrate binding mechanism. All structural diagrams presented in this paper were prepared with PyMOL (Schrodinge, LLC, The PyMOL Molecular Graphics System, version 2.5.2).

Both the N2 and N3 domains are of so-called described previously Dev-IgG like fold (**Figure 3F**) (8). Commonly, there are some variations on the edge strands (D and D1 side) of the fold. In *S. aureus* SdrD, the D strand is missing in the N2 domain, while the fold of N3 domain is more typical, with only a short A^0^ strand antiparallel to A strand (**Figure 3C and F**). Besides a single peptide linker between the two N domains, contact between them is modest with buried surface area (BSA) ranging from 960 –1,030 Å^2^ in the three structures, including at least three hydrogen bonds (**Figure S5**). As a result of packing of the two N domains through their loops, largely the loop between E and F strands (E_F loop) of N2 and the long D_D1 loop of N3 (**Figure 3C**), the N2N3 conformation seems to be quite stable, with relative rotation between the two domains limited to about 4° among the three structures (**Figure 4A**). The residue P393 on the short link between N2 and N3 likely helps limit relative motion between these two domains (**Figure S5**).

**Figure 4.**
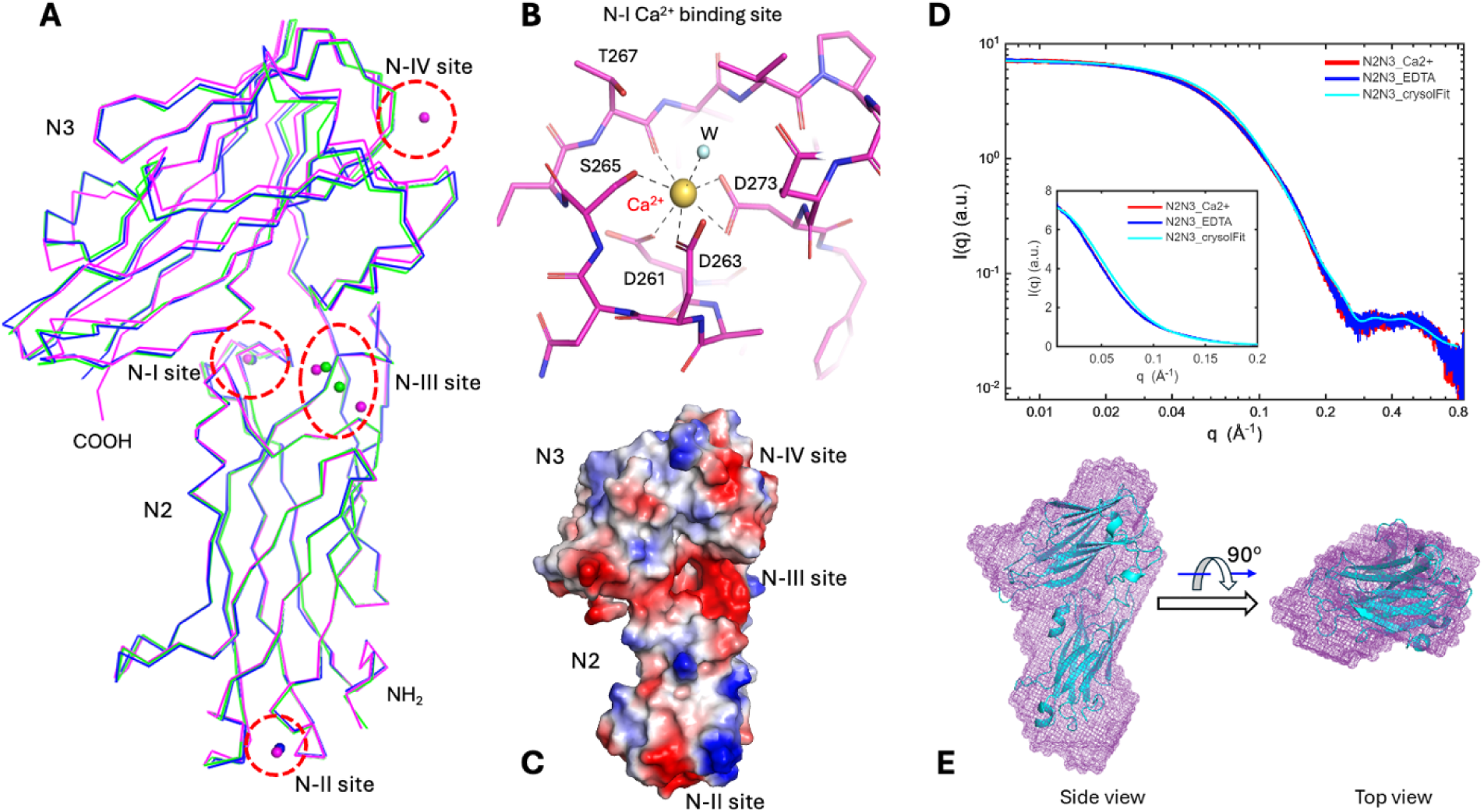
N2 and N3 domains and their Ca^2+^-binding sites. (A) The rigidity of N2 and N3 domains and their relative orientation. The N2 domains from N2N3B1 (in blue) and N2N3B1B2 (in green) structures are superimposed to the N2 domain of N2N3 structure (in magenta) by using least square structural alignments. The RMSD (root-mean-square deviation) values are less than 0.5 Å. Limited variations (< 4°) in the relative rotation of N3 in respective to N2 were observed. Four Ca^2+^-binding sites (N-I, –II, –III and –IV) are marked in dashed oval/circles. (B) The N-I Ca^2+^-binding site is like a canonic EF-hand Ca^2+^-binding site but without a helix. The diagram is drawn in stick format to detail hydrogen bonds. (C) An electrostatic potential surface representative of the N2N3 domain (blue, positive; red, negative). The weak N-II, –III and –IV Ca^2+^-binding sites in the view are associated with negative potential (red) patches on the molecular surface. The conserved N-I site is out of view. (D) Superimposed SAXS profiles of N2N3 in protein buffer (red curve), in the protein buffer with 10 mM EDTA (blue), and fitting with the crystal structure (cyan) using the program CRYSOL. Inset: the three SAXS profiles displayed in linear scales. The SAXS profiles for samples in the absence or in the presence of EDTA are virtually identical. (E) Side and top views of the SAXS envelope derived from SAXS data in the presence of Ca^2+^. The SAXS envelope (in mesh mode and purple color) was overlaid with the N2N3 crystal structure (in cartoon mode and cyan color).

On the A-B loop of N2, there is a conserved EF hand-like Ca^2+^-binding site (N-I, **Figure 4A-B**), formed by D261, D263, S265, T267 (O atom) and D273 (bidentate); and completed by a water molecule at one axial position of the pentagonal bipyramidal configuration. A non-conserved Ca^2+^-binding site (N-II) is located at the bottom of N2 domain, which is shallow, with the metal ion chelated, mostly by carbonyl groups and surface water(s). Additionally, across the interface between two N domains (N-III) there is a cluster of negatively charged residues, including D299, D363 and D365 from N2 domain, and D481 and E410 from N3 domain. These residues form Ca^2+^-binding site(s) at different locations and in different numbers in three N2N3-containing structures. The structures of both the N-II and –III sites indicate they are opportunistic Ca^2+^-binding sites. There is only one such weak Ca^2+^-binding site found in N3 domain in one structure (N-IV in **Figure 4A** and **4C**). The three weak Ca^2+^-binding sites are generally associated with negative surface patches in electrostatic potential surface representation (**Figure 4C**). It is unclear if the opportunistic Ca^2+^-binding sites have any functional impact on SdrD. The N-III site lays between two N domains and sometimes bridges them. The N-III site could be a factor in regulation of the relationship between the two N domains, corresponding to different relative rotation angles mentioned above.

Considering the critical roles the two N domains play in the widely accepted DLL peptide substrate-binding mechanism for SdrD and other family members, small-angle X-ray scattering (SAXS) experiments were performed to investigate the conformation and potential conformational changes of the N domains in the presence and absence of Ca^2+^. Experimental data show the N2N3 solution structure is consistent with its crystal structure (**Figure 4D**). The calculated SAXS profile from the crystal structure resembles the Ca^2+^ experimental SAXS profile across the entire range, with a relatively large chi2 value of 7.2. The main discrepancy appears in the q range of 0.04 – 0.1 Å^-1^ with calculated SAXS intensities being systematically higher than experimental values (**Figure 4D**), indicating that the solution-state N2N3 is slightly larger than its crystal form. This observation is further supported by the 3D SAXS molecular envelope and PDDF analysis. The solution-state N2N3 molecular envelope derived from SAXS data exhibits a shape similar to that of the crystal structure, but is slightly larger in solution state (**Figure 4E**); the PDDF analysis suggests the solution-state N2N3 is ∼6 Å larger than its crystal form, potentially due to an hydration layer and a looser packing of the molecule in solution (**Figure S4C**).

When N2N3 was treated with 10 mM EDTA, there were no significant conformational changes after removal of Ca^2+^ from N2N3, evidenced by virtually identical SAXS profiles and R_g_ values under the two conditions (**Figure 4D** and **S4B**). This suggests that the three Ca^2+^-binding sites, N-II, N-III, and N-IV, are likely opportunistic, and that the EF-hand-like Ca^2+^-binding site N-I, located on the A–B loop of N2, has only a minor impact on overall folding of N2N3, even though this loop is expected to become more flexible in the absence of Ca^2+^.

### B domains

There are five B domains in *S. aureus* SdrD, with conserved structural folds and Ca^2+^-binding sites (**Figure 5A**). Besides the N2N3B1 construct, N2N3B1B2 construct was produced to better understand the connection between B domains. The B domain fold is of cnaB type (or IgG-rev type), in which the edge F strand can shift between two opposite ß-sheets. In *S. aureus* SdrD, the DAG strands are on one side and the CBEF strands on the other (**Figure 5B**). There is a short strand on the edge by G strand, labeled as the F^0^ strand. There are three conserved Ca^2+^ binding sites on each B domain. Taking the B1 domain as an example, the B1-I site is an EF-hand like Ca^2+^-binding site located in the A-B loop, immediately after the A strand. This is like the N-I site on the N2 domain described above (**Figure 4A** and **B**). However, both B1-IIa and B1-IIb sites are solvent accessible. The Ca^2+^ at each of the two sites is chelated by two sidechains and one mainchain carbonyl group and completes its coordination with three water molecules. The two sites are adjacent to on another and bridged by a highly conserved bidentate aspartic acid (on G strand) from B1-IIa site and a water molecule from the B1-IIb site through a hydrogen bond. Based on formation of these three Ca^2+^ binding sites, the B1-I site is likely the high affinity (Class I) site, while the B1-IIa and –IIb sites are low affinity (Class II) sites of the B1 domain (64). Each of other four B domains of SdrD similarly have one high affinity and two low affinity Ca^2+^ binding sites. Interestingly, all three B domain Ca^2+^ binding sites are largely on one side (DAGF^0^) of two ß-sheets (**Figure 5C**). B1-I and B1-IIb are apparently critical in fixing conformations of the long A-B loop and F^0^-F loops, respectively. B1-IIa is on the outside of the G strand, nearly in the middle of a negatively charged shallow groove. The Ca^2+^ ion on this site is completely exposed, seemingly accessible to a potential ligand.

**Figure 5.**
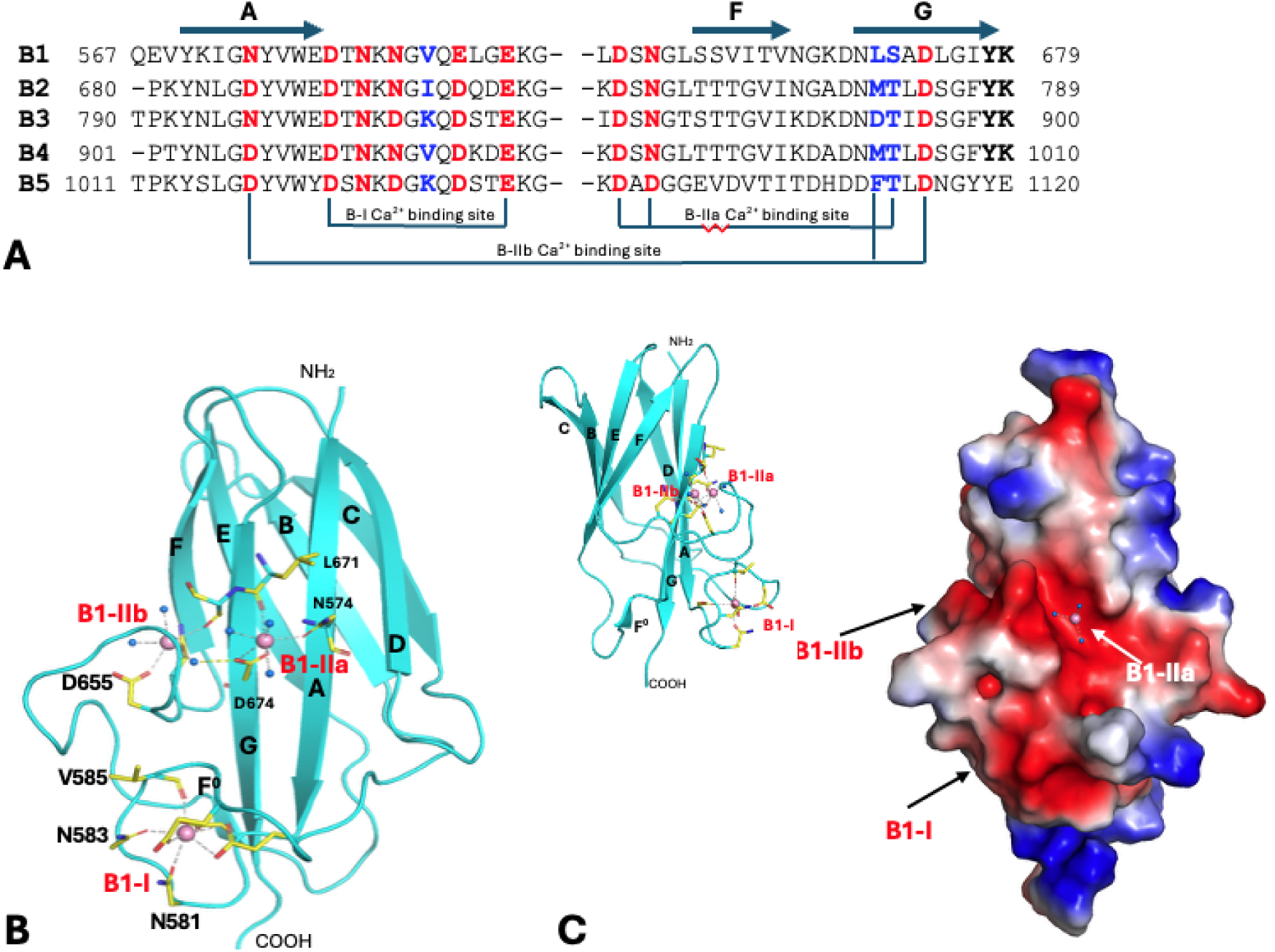
B domains and their Ca^2+^-binding sites. (A) A partial sequence alignment of five consecutive B domains of *S. aureus* SdrD illustrating the conservation of three Ca^2+^-binding sites, B-I, B-IIa and B-IIb. See text for naming of the Ca^2+^-binding sites. The B-I site is an EF-hand like Ca^2+^-binding site right after A strand. Residues that use their sidechains for Ca^2+^ coordination are highlighted in red. Residues that use their mainchain carbonyl groups for Ca^2+^-binding are marked in blue. Protein atoms contribute to the B-IIa and B-IIb Ca^2+^-binding sites. The conserved motif YK at the end of B1-B4 domains are highlighted in bold. (B) Ribbon-diagram of B1 structure, a representation of five B domains. All ß-strands of B1 domain are labeled. An extra short strand preceding the F strand and antiparallel to the G strand is labeled as the F^0^ strand, for convenience. Residues contributing to Ca^2+^-binding are shown in stick format. Only some are labeled for clarity of presentation. Both B1-IIa and B1-IIb sites are solvent accessible, requiring two and three water molecules for complete chelation for Ca^2+^. In the top-right corner, a small diagram of the B1-I domain viewed from one side shows that three Ca^2+^-binding sites are largely on one side of the domain, the GAD ß-sheet. (C) An electrostatic potential surface representative of B1 domain. The B1-IIb Ca^2+^-binding site is within an extensive negative potential (red) patch and seemingly ready for a ligand to access. The small ribbon diagram on the left top shows that the three Ca^2+^ bindings sites on B domain are on one side of the ß-sheet.

Prior studies report that B domains are unfolded in the absence of Ca^2+^ (64, 65), and propose that they act as spacers and/or springs between ligand-binding N2N3 domain and bacterial membranes, positioning and orienting the N2N3 domains towards targeting molecule(s) in the presence of a Ca^2+^ gradient. To assess the solution property of B domain, SAXS experiments were performed with a single B1 domain. In protein buffer, the solution structure is predicted to be consistent with its crystal structure, as indicated by the low chi2 value (0.58) between the experimental SAXS data and the profile calculated from the crystal structure (**Figure 6A**), as well as the excellent agreement between the crystal structure and reconstructed SAXS envelope (**Figure 6B**). After the B1 domain was treated with 10 mM EDTA in protein buffer, there was a significant change of SAXS profile (**Figure 6A**). The radius of gyration (Rg) increased from 16.6 to 20.2 Å, and the PDDF analysis indicated the maximum dimension (Dmax) of B1 to have increased by up to ∼20 Å after removing Ca^2+^ (**Figure S6 B and C**). However, the partially Bell-shaped Kratky plot (**Figure 6A**) suggested that the B1 domain is not completely unfolded upon Ca^2+^ removal. Consistently, the reconstrued SAXS molecular envelope (**Figure 6B**), indicated that the core of the B1 domain remains folded after Ca^2+^ removal, whereas the Ca^2+^ binding loops could loosen and become more elongated, as evidenced by the peak position shift in the dimensionless Kratky plot (**Figure 6A**, inset).

**Figure 6.**
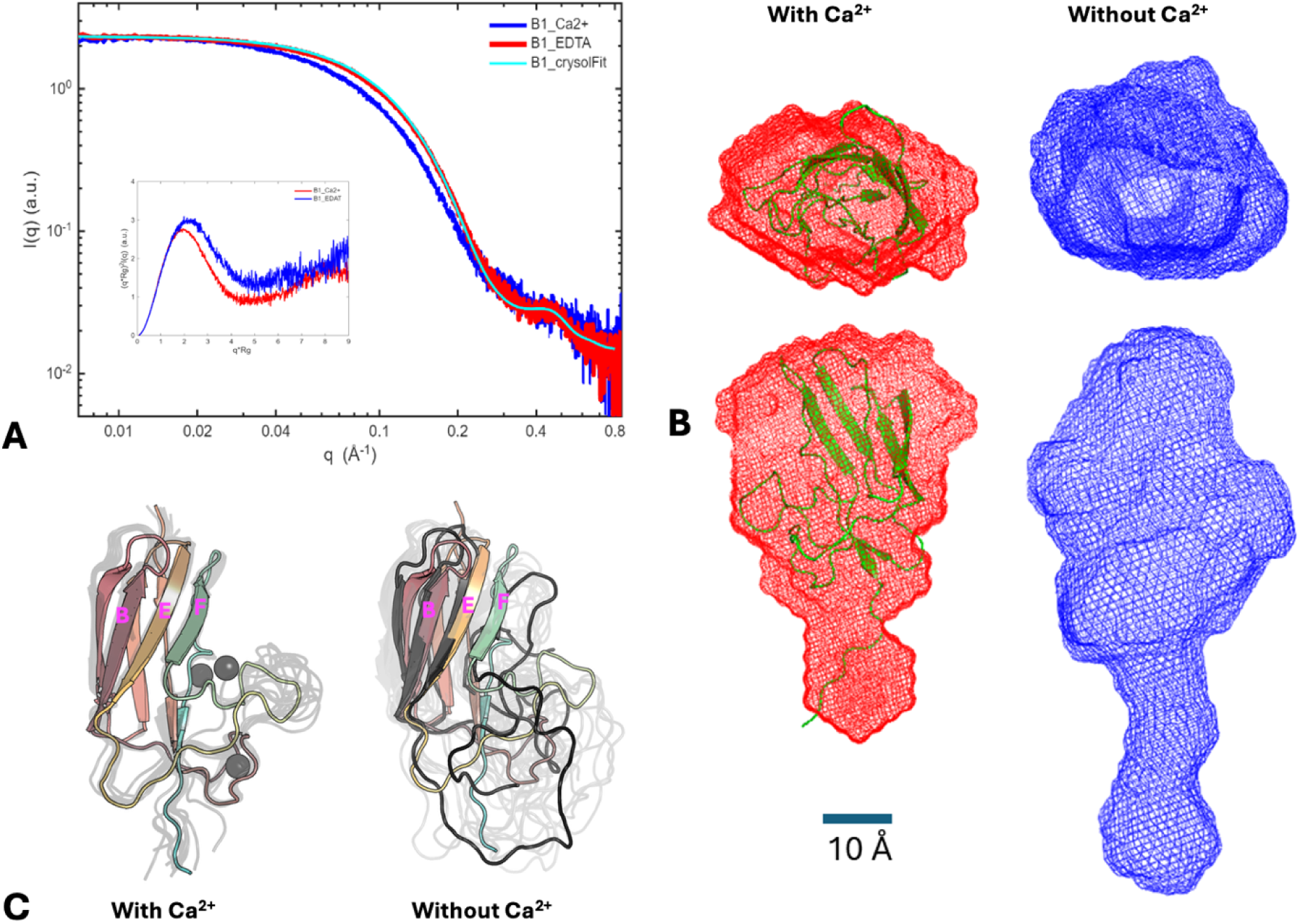
SAXS and MD data on B1 domain. (A) Superimposed SAXS profiles of B1 in protein buffer with Ca^2+^ (red curve), in protein buffer with 10 mM EDTA added (blue), and fitting with the crystal structure (cyan). The goodness-of-fit (chi2) against the red curve is 0.58. Inset: dimensionless Kratky plots for the SAXS data. Colors were coded as with the SAXS data. The partially Bell-shaped blue Kratky plot suggested the B1 domain was not completely unfolded upon Ca^2+^ removal. The blue Kratky peak shifts to a higher value, suggesting B1 becomes more elongate after removing Ca^2+^. (B) Top and side views of the most-probable SAXS molecular envelopes for B1 reconstructed from SAXS data under the two conditions. Color coding is as in A-C. The crystal structure was displayed in green color and in cartoon mode. (C) Overlay of the conformational ensembles for B1 with Ca^2+^ (left) and B1 without Ca^2+^ (right). The crystal structure is provided with rainbow colors in the background. Representative structures from clustering based on the RMSD are overlayed in gray. Ca^2+^ ions are show as grey spheres. A representative conformation is shown in black for B1 without Ca^2+^.

To further verify this observation, MD simulations were performed. MD simulations carried out at 300 K displayed minimal differences in the conformational ensemble, suggesting preserved stability at 300 K without the presence of Ca^2+^ (**Figure S7**). Higher temperature simulations carried out at 450 K revealed partial unfolding of the B1 domain that would otherwise be stabilized by the Ca^2+^ (**Figure 6C**). In the absence of Ca^2+^, the unstructured B1 loop regions became flexible, increasing structural heterogeneity for the B1 domain, and resulting in a more elongated shape. These observations are consistent with the SAXS findings, where addition of EDTA resulted in an increase in the *R_g_*; however, the B1 domain remained compact, as evidenced by the Kratky plot (**Figure 6A**). Calculation of the theoretical SAXS for the B1 domain with and without Ca^2+^ aligned with the experimentally observed SAXS profiles, confirming that the tertiary structure of the B1 domain is largely preserved upon removal of Ca^2+^ (**Figure S7**).

These results were unsurprising and consistent with the structural observation that Ca^2+^ binding sites on B domain are either associated with loop fixation (B-I and B-IIb sites) or on the ß-sheet surface (B-IIa site), but not directly involved in the folding of B domain core.

### Association between N2 and B1 domains

The unique triangular conformation formed by the N2, N3 and B1 domains largely depends on contact between N2 and B1, while the linker between N3 and B1 is flexible (**Figure 3D** and **E**). The similar trio-domains arrangement has been observed in a homolog *S. aureus* SdrD structure (29), and to some extent in the UafA structure (36), in which the B1 domain contact to the N2 domain seems unrestricted. However, it was unclear whether the seemingly conserved conformation had any significance or was simply a result of molecular packing in crystals. More importantly, such a triangular conformation, if stable, would not compromise a potential DLL mechanism proposed for DsrG and others, in which the B1 domain needs to move away from N2 domain readily so that the linker between N3 and B1 can lock the bounded ligand peptide aside of the N3 domain and latch into the N2 domain. Therefore, the nature of N2-B1 interface observed in the structure is a critical factor to applicability of the DLL mechanism in SdrD.

The center of the N2-B1 interface on the B1 domain is the solvent exposed B1-IIa Ca^2+^-binding site, as described and emphasized above, while the interface center on N2 domain is the E324 on the protrude D^0^-D1 loop (**Figure 7A**). A water molecule bridges the metal ion and glutamic acid, while E324 forms an additional hydrogen bond to N547 (from B1), also coordinated to Ca^2+^. There are two more hydrogen bonds between N2 and B1 and several hydrophobic and van deer Waals contacts across the interface, with a BSA value of about 842 Å^2^. The BSA value does not appear large enough, however, to guarantee a stable protein-protein interface (66). The B1-IIa Ca^2+^-binding site likely provides additional support for the N2-B1 interaction. The key feature to the interaction between the two domains may be simply described as docking of a protruded glutamic acid onto a Ca^2+^-binding site.

**Figure 7.**
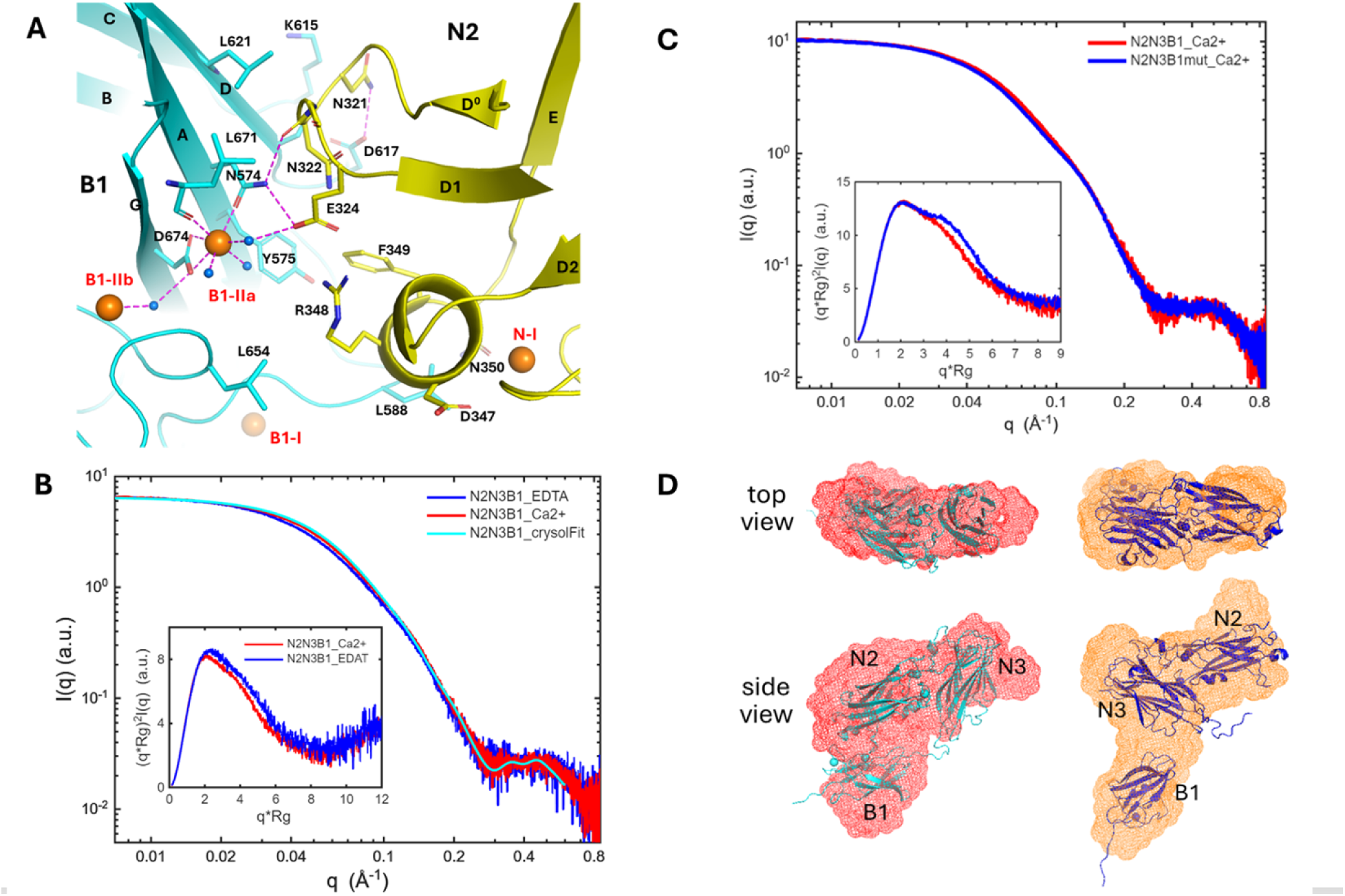
Association between N2 and B1. (A) A ribbon diagram of a zoom-in view of the interface between N2 (in yellow) and B1 (in cyan). Residues involved in the interaction from two domains are drawn in stick format. Coordination bonds to Ca^2+^ ion and hydrogen bonds are drawn in magenta dash lines. On the B1 domain, the Bb Ca^2+^-binding site is the interaction center. On the N2 domain, the protruded loop between D^0^ and D1 strands is the primary structural element participating in the interaction. The three residues on the loop, N321, N322 and E324, especially E324, play the key role on N2 domain side. (B) Superimposed SAXS profiles of N2N3B1 in protein buffer with Ca^2+^ (red), with EDTA (blue), and fitting with the crystal structure (cyan). Inset: corresponding dimensionless Kratky plots for the SAXS profiles in (B). (C) Superimposed SAXS profiles of N2N3B1 (red) and N2N3B1mut (blue) in protein buffer with Ca^2+^. Inset: Corresponding dimensionless Kratky plots for SAXS profiles. (D) Side views of the refined SAXS molecular envelope for N2N3B1 (red) and N2N3B1mut (orange) reconstructed for data collected in protein buffer with Ca^2+^. Left: the N2N3B1 crystal structure was overlaid with N2N3B1 SAXS envelope and displayed in cyan color and in cartoon mode. Right: the N2N3B1 crystal structure was divided into two parts, i.e., N2N3 and B1, rearranged and overlaid with N2N3B1mut SAXS envelope.

The SAXS experiment with N2N3B1 shows that the structure of the tri-domain construct is a stable entity in solution. The SAXS profile fits well to N2N3B1 crystal structure (**Figure 7B**), with a decent chi2 value of 2.2, better than for the N2N3 construct. Consistently, the reconstructed SAXS molecular envelope shows good agreement with the crystal structure (**Figure 7D**). Upon EDTA treatment, only modest changes are observed in SAXS data. Accordingly, both the R_g_ value and Dmax increase after removing Ca^2+^ (**Figure S8B and C**). The Kratky plots exhibit similar shape under both conditions, with only a slight shift in peak position (**Figure 7B, inset**), indicating that N2N3B1 retains the same overall conformation but becomes slightly more elongated due to loosening of B1 domain upon Ca^2+^ removal.

To probe the interface between N2 and B1, a N574A mutation was introduced on B1; this removes two hydrogen bonds at the interface and probably disrupts the B1-IIa Ca^2+^-binding site (**Figure 7A**). The mutation leads to noticeable changes in the SAXS profiles, particularly in the q range of 0.03 – 0.15 Å^-1^, reflecting alternations in overall size and potentially conformation. Guinier analysis shows that the mutation increases the R_g_ value from 31.2 to 32.8 Å (**Figure S9A** and **S9B**); the PDDF analysis shows a molecular size increase of 10-20 Å for the mutant construct (**Figure S8C and S9C**). In addition, there are two kinks in the mutant SAXS profile (i.e., at ∼0.04 and ∼0.15 Å^-1^) and a second peak in the Kratky plot of 4-6 at q*R_g_, indicative of a branched or bent conformation (**Figure 7C**, inset). Consistently, the reconstructed SAXS molecular envelope of the mutant adopts an “L” conformation, suggesting a rearrangement of the B1 domain with respect to the N2-N3 domains upon disruption of the two interdomain hydrogen bonds between N2 and B1. It is speculated that the B1 domain moves away from N2 domain and that the N2N3B1 mutant adopts a “L” shape, like the orange conformation in **Figure 7D**. This “L”-shaped conformation successfully reproduces the experimental SAXS and Kratky features observed for the mutant (**Figure S9**).

EDTA treatment of the N2N3B1 mutant produces minimal changes in SAXS profiles, particularly at q <0.2 Å⁻¹, indicating no significant global structural changes upon Ca²⁺ removal for the mutant.

### B-B domain connection

Structural determination of the N2N3B1B2 construct enables understanding of the connection between B domains, which have been regarded as only spacers to project functional N domains. Linkers between consecutive B domains are short and rigid. Proline alone constitutes the linker between B1–B2 and B3–B4, whereas the B2–B3 and B4–B5 linkers are Thr–Pro dipeptides. The C-terminal sequence of each B domain is highly conserved and includes a Tyr–Lys motif (**Figure 5A**) preceding the linker to the next B domain. In B1, the tyrosine of this motif (Y678) forms a CH–π stacking interaction with K582, a conserved lysine from the EF-hand–like Ba Ca^2+^-binding site on the A–B loop (**Figures 5A** and **8A**). The lysine of the motif (K679) forms a second CH–π stacking interaction with a conserved tyrosine (Y641) immediately preceding the short F^0^ strand (**Figure 8A**). The two conserved CH–π stackings reinforce packing at the bottom of each B domain. Notably, each B domain typically begins, immediately after a B–B linker, with a Lys–Tyr dipeptide. In B2, the lysine (K681) forms a salt bridge with D737 on the D–E loop; in B3–B5, the residues at the equivalent position are Glu, Asn, and Asp, respectively, potentially forming similar salt bridges or hydrogen bonds with the N-terminal lysine of the B domain. The tyrosine of the Lys–Tyr motif packs against the tip of the F–G loop. Together, these interactions compact the top of the B domain. Given the short, relatively rigid linkers between B domains, their relative motion is expected to be limited. This structural observation is consistent with an earlier analysis based on gel filtration data, suggesting that the shape of five B domains together is a rigid rod (64).

**Figure 8.**
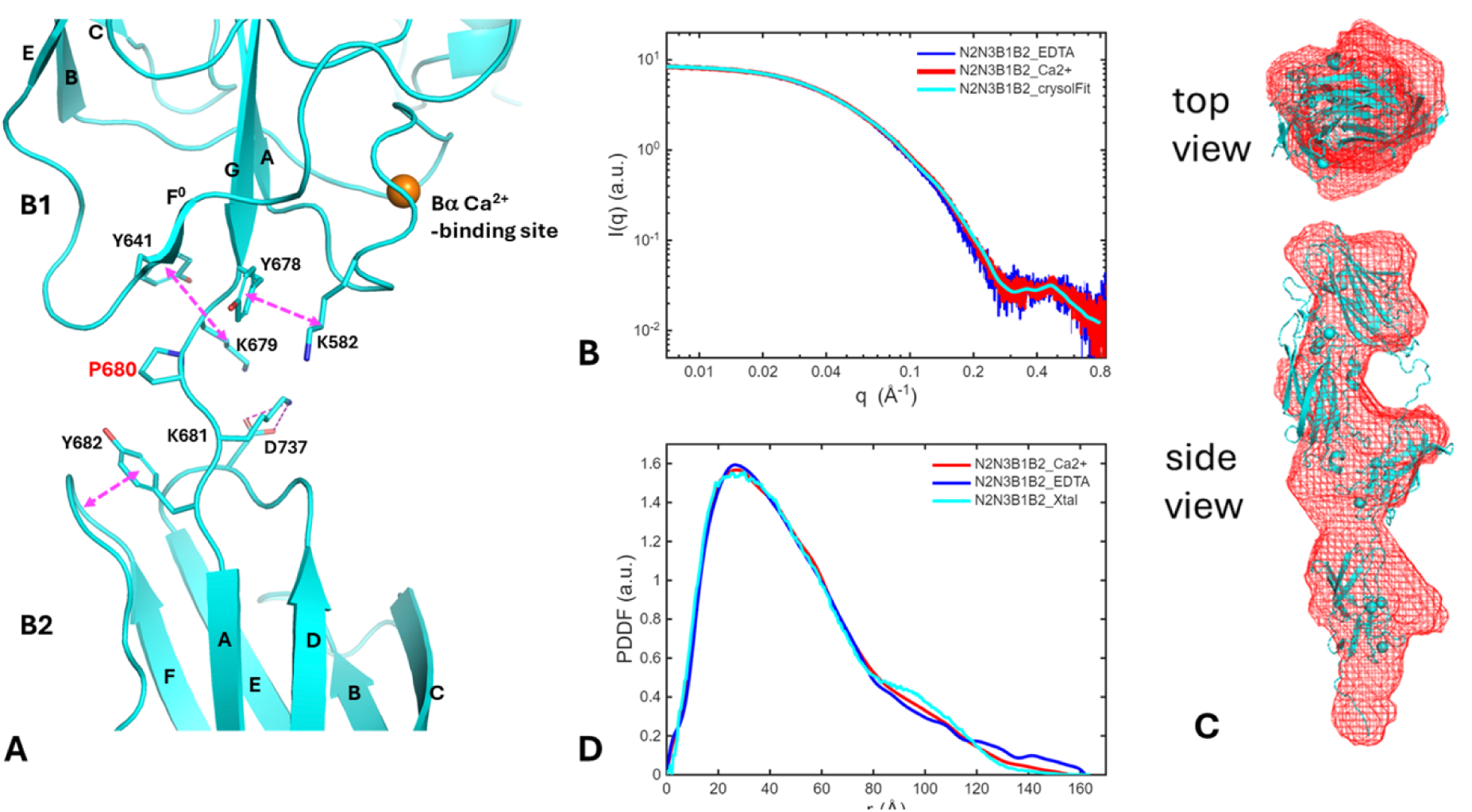
B-B domain connection. (A) The connection between B1 and B2 domains is shown, considering the availability of structural information and highly conserved residues involved in the link and contact between any two B domains. To illustrate the interface between the B1 and B2 domains, the bottom half of B1 and the top half B2 domains are displayed. CH–π stackings and the interaction between the aromatic ring and peptide backbone are marked with magenta double arrow dash lines. Salt bridges are drawn in simple magenta dash lines. (B) Superimposed SAXS profiles of N2N3B1B2 in protein buffer with Ca^2+^ (red curve), in protein buffer with 10 mM EDTA added (blue), and fitting with the crystal structure (cyan), respectively. Goodness-of-fit (chi2) of CRYSOL fitting with the crystal structure against the red curve is 0.44. (C) The top and side views of the most-probable SAXS molecular envelope for N2N3B1B2 reconstructed for the red SAXS profile in (B). The N2N3B1B2 crystal structure was displayed in cyan color and in cartoon mode and superimposed with the SAXS molecular envelope. (D) PDDF profiles for N2N3B1B2. Red, experimental PDDF calculated from the red SAXS data in Figure 6B using GNOM; blue, experimental PDDF calculated from blue SAXS data in Figure 6B; cyan, theoretical PDDF calculated from the crystal structure using SolX3. To highlight the difference among the PDDFs, the curves were normalized by their respective areas.

Solution SAXS data were collected for the N2N3B1B2 construct in protein buffer with Ca^2+^ and in buffer with added 10 mM EDTA (**Figure 8B**). The crystal structure of N2N3B1B2 agrees with the solution SAXS data in the presence of Ca^2+^, with a super chi2 value of 0.44. Consistently, the reconstructed SAXS molecular envelope exhibits a rod-like, elongated conformation, aligning with the crystal structure (**Figure 8C**). This suggests that the crystal structure provides a reliable model for the solution conformation of this construct.

Removal of Ca^2+^ with EDTA did not induce significant changes in the SAXS profile. The R_g_ value increased modestly from 37.5 to 38.3 Å (**Figure 8D**). The PDDF profile amplitude slightly decreased in the distance range of 75-115 Å and increased at distances of 115-160 Å, accompanied by an extension of the D_max_by approximately 5-10 Å (**Figure 8D**). The distance range of 80-120 Å corresponds to the correlation between the N3 and B2 domains. These changes in molecular size and PDDF most likely arise from loosening of loops in B domains upon Ca^2+^ removal. Nevertheless, the overall rod-like, elongated conformation is expected to be preserved.

Solution SAXS data were also collected for a longer construct, N2N3B1B2B3 under the two conditions. An atomic model of this construct generated using AlphaFold3 (AF3) (67), closely resembles the crystal structure of N2N3B1B2, with the addition of one extra B3 domain. This AF3 model fits the Ca²⁺ solution SAXS data well, with an excellent chi2 value of 0.69 (**Figure 9A**). In the PDDF profile for the Ca^2+^ solution condition (red, **Figure 9B**), two broad peaks are observed in the distance range of 80–170 Å, corresponding to interdomain distances between the N and B domains. The peak positions agree with those derived from the theoretical PDDF (cyan, **Figure 9B**). These results indicate that the AF3 model provides a good representation of the solution conformation of the construct in the presence of Ca²⁺.

**Figure 9.**
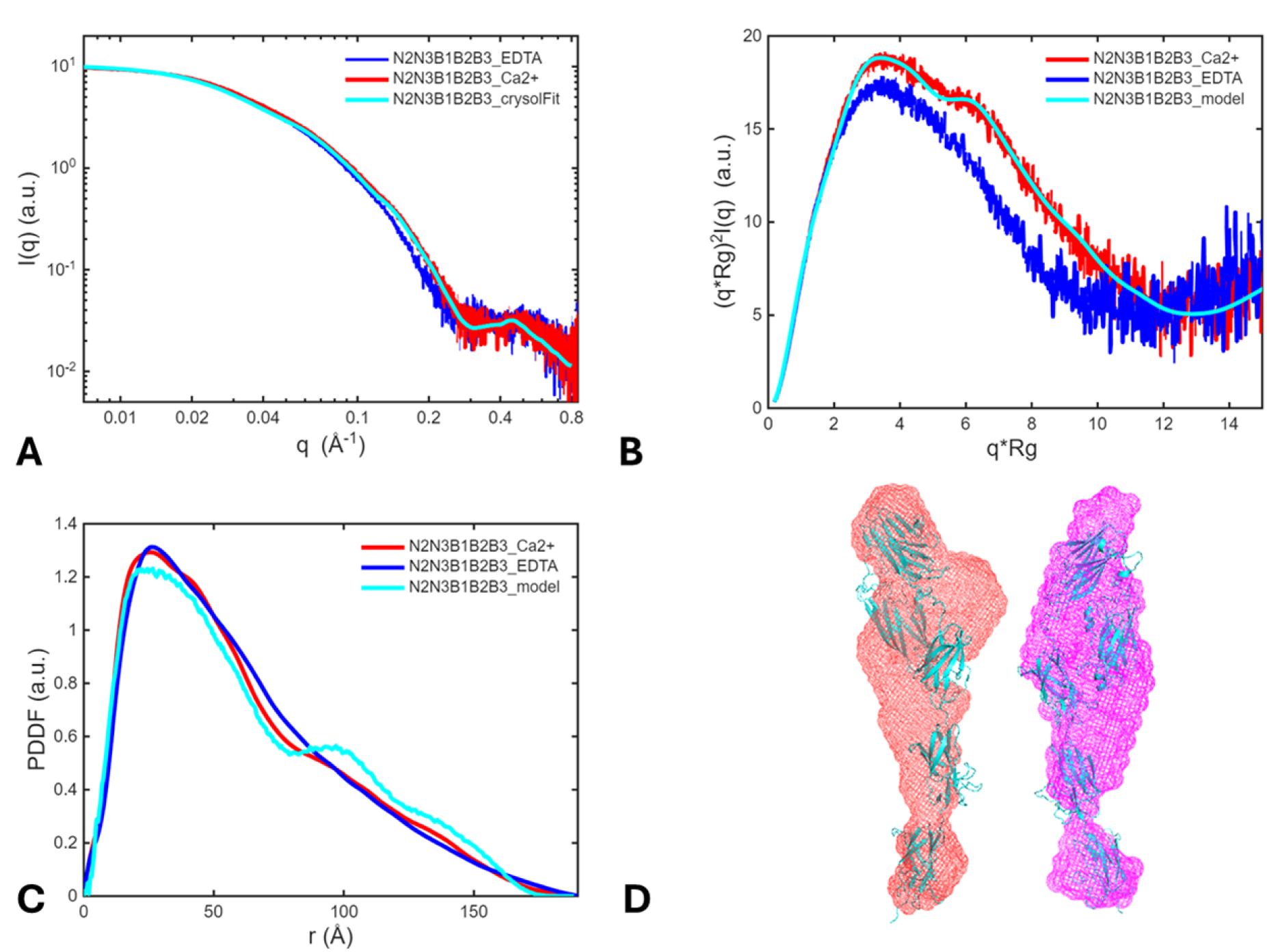
SAXS results for protein N2N3B1B2B3. (A) Superimposed SAXS profiles of N2N3B1B2B3 in protein buffer (red curve), in protein buffer with 10 mM EDTA added (blue), and the fitting with the alphfold3 (AF3) predicted atomic structure (cyan). The goodness-of-fit (chi2) of CRYSOL fitting with the AF3 atomic structure against the red curve is 0.69. (B) Dimensionless Kratky plots for N2N3B1B2B3 under both solution conditions, and the crystal structure fitting. (C) PDDF profiles for N2N3B1B2B3. Red, calculated from experimental SAXS data (red curve in Figure 7A) using GNOM; blue, calculated from blue SAXS data in Figure 7A; cyan, calculated from the AF3 structure using SolX3. (D) Side views of the SAXS molecular envelope for N2N3B1B2B3 reconstructed for data collected with Ca^2+^ (left, red mesh, most-probable model) and EDTA (right, magenta mesh, refined model), superimposed with the AF3 model (cyan color).

As with the N2N3B1B2 construct, EDTA treatment induced only minor changes in N2N3B1B2B3, including a modest increase in R_g_ from 45.1 to 45.6 Å and attenuation of the PDDF peaks in the 80–170 Å range, likely reflecting loosening of loops in the B domains upon Ca²⁺ removal. Nevertheless, the construct adopts a rod-like, elongated conformation under both conditions, as evidenced by the reconstructed SAXS molecular envelopes (**Figure 9C**).

In summary, solution SAXS measurements confirm that the crystallographic and AF3 models provide good representations of the solution-state conformations for each construct in the presence of Ca²⁺. This agreement is particularly strong for B1-containing and longer constructs, N2N3B1B2 and N2N3B1B2B3, as evidenced by the excellent fit values obtained using these atomic structures. The largest discrepancy in structural fitting, as judged by the chi2 value, was with the N2N3 construct, whereas the chi2 value was reduced for N2N3B1. This improvement may result from two possible factors. First, binding of B1 to the N2 domain may render the N2N3 module more compact and closer to its crystallographic conformation. Second, excellent agreement for the B1 domain and the correct overall arrangement in N2N3B1 reduce the relative contribution of deviations in the N2N3 region to the overall chi2 value.

The Kratky plot provides a rapid assessment of molecular conformation. In the dimensionless Kratky plots for constructs ranging from N2N3 to N2N3B1B2B3 (**Figure 10A** and **S11**), the peak position progressively shifts to higher values, indicating that the constructs become increasingly elongated as additional B domains are added. In addition, a shoulder appears on the Kratky peak and becomes more pronounced for the longer constructs. This feature likely arises from a slightly bent conformation between the N2N3 module and the B-domain tract. With EDTA treatment, attenuation of this shoulder is observed for N2N3B1B2 (**Figure S10**) and N2N3B1B2B3 (**Figure 9B**), suggesting modest rearrangement between N2N3 and the ß-tract upon Ca²⁺ removal, resulting in a less bent overall conformation. In contrast, disruption of hydrogen bonds between B1 and N2 by mutation abolishes their interaction, as observed for the N2N3B1 mutant, leading to a large rearrangement of the B domains relative to the N2 domain.

**Figure 10.**
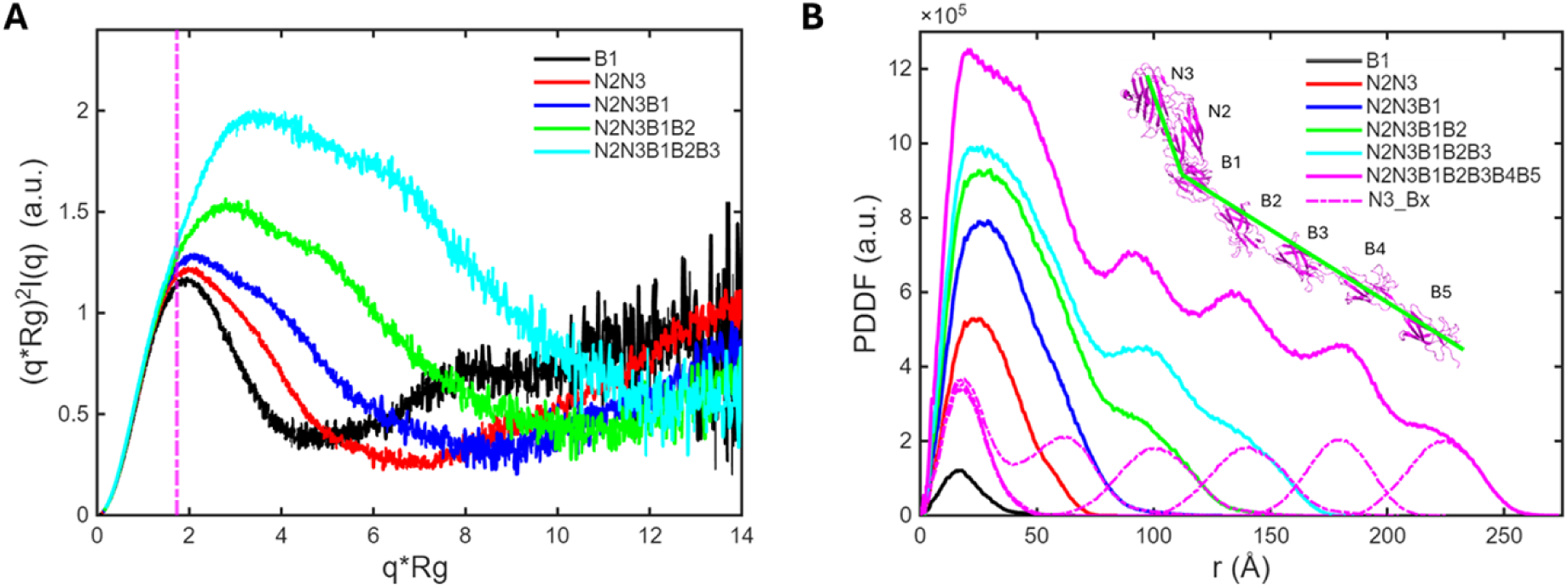
SAXS results comparison for all constructs. (A) Dimensionless Kratky plots for experimental SAXS data. The magenta vertical line is q*Rg=√3, where the peak of spherical particle appears. The higher the aspect ratio of length to cross-section, the further the peak position away from √3. (C) Theoretical PDDFs of all constructs calculated from atomic structures. Among thoe atomic structures, B1, N2N3, N2N3B1 and N2N3B1B2 are crystal structures, and N2N3B1B2B3 and N2N3B1B2B3B4B5 are AF3 models. N2N3B1B2B3B4B5 is displayed in the inset with labels of N and B domains. The dash line profiles in magenta are five PDDFs for the ensemble structure consisting of N3 and one individual B domains, i.e., N3_B1, N3_B2, N3_B3, N3_B4, and N3_B5, respectively. They consist of two peaks, one short distance peak centering at ∼18 Å, and one far distance peak centering from 62 to 224 Å, depending on the location of B domain. The first peak reflects the intra-domain pair-wise distance within N3 or B domains, while the second peak describes the inter-domain pair-wise distance correlations between N3 and B domains. The peak position of the second peak reflects the inter-domain center-to-center distance. The green lines on the AF3 model highlight the bent conformation for this structure.

SAXS PDDF profiles provide additional insight into the molecular conformation. Simulations indicate that PDDF peaks at longer distances are dominated by distance correlations between N3 and B domains (**Figure 10B**). These distinct peaks arise from a bead-on-string architecture with well-defined boundaries between domains. Consistent with this model, experimental PDDFs for the longer constructs, N2N3B1B2 and N2N3B1B2B3, display pronounced long-distance peaks under Ca²⁺ conditions. Following EDTA treatment, the peaks are attenuated, most likely due to loosening of the Ca²⁺-binding loops, which transforms the ß-tract from a well-defined bead-on-string arrangement into a more fuzzy rod–like conformation.

Notably, the overall sizes of most constructs change only modestly with Ca²⁺ removal, suggesting that linkers between B domains are relatively rigid and that each B domain maintains its size and spacer function. Taken together, these observations indicate that the ß-tract adopts a bead-on–rigid-string conformation.

### Exploration of SdrD-1 ligand binding

Desmoglein-1 (DSG-1), a cell adhesion protein found in the outer layers of the skin, has been identified as a potential host binding partner for SdrD and possibly other *S. aureus* surface adhesion molecules (12). This bacterium–host molecular interaction plays a role in promoting *S. aureus* adhesion to host cells (12). A recent study demonstrates that the SdrD-DSG-1 interaction is exceptionally strong and displays characteristic catch-bond features (18). The study also identified two potential SdrD binding sites on DSG-1, P_dist_ and P_prox_. The P_dist_ is largelt the N-terminal peptide before the first domain of DSG-1, which is a type of desmosomal cadherin (18). In consideration of the common cadherin structures, the N-terminal peptide was thought to be a likely candidate to participate in binding to an edge strand of SdrD N3 domain based on DLL binding mechanism. Thermal shift experiments were performed to assess P_dist_ binding to the various SdrD constructs mentioned above; however, results were inconclusive (data not shown). The peptide was also used in co-crystallization trials with selected SdrD constructs (N2N3 and N2N3B1). However, all crystals obtained were SdrD alone, with no peptide co-crystals observed.

## DISCUSSION

Here, the structure of *S. aureus* JH1 SdrD was determined using X-ray crystallography, complemented by solution small-angle X-ray scattering and MD simulations. Together, the data reveals structural features – including the architecture of individual domains, their interdomain organization, and the behavior of the calcium-binding B domain upon calcium depletion – that prompt a re-evaluation of prevailing models for the Sdr subfamily, including assumptions about domain flexibility and possible ligand-binding mechanisms. These structural findings are further contextualized by population-level genomic analyses, which demonstrate that *sdrD* exists as three sequence subtypes across the *S. aureus* population and that the JH1 subtype is the predominant variant among clinical PJI isolates, making the JH1 SdrD structure presented here of direct relevance to understanding *S. aureus* pathogenesis in PJI.

Phylogenetic analyses of the *sdrCDE* locus reveal that *sdrD* is not a monolithic gene but instead exists as three sequence subtypes across the *S. aureus* population, represented by USA300, JH1, and TCH60 reference strains. This diversity was initially uncovered by observing that BLAT-based genomic searches using USA300 and JH1 as independent queries returned non-overlapping sequence sets of differing lengths, suggesting that the subtypes are sufficiently diverged to escape cross-detection. Phylogenetic reconstruction using both reference strains and a large public genome library confirmed three discrete *sdrD* nodes at both the nucleotide and amino acid levels, with each node forming a well-supported cluster, distinct from sdrC and sdrE. To contextualize these findings in a clinically relevant setting, the analysis was extended to a collection of 156 *S. aureus* isolates derived from patients with PJI. Consistent with prior reports implicating *sdrD* in bone and implant-associated disease, *sdrD* was detectable in a substantial proportion of PJI isolates, though prevalence varied depending on which reference sequence was used as a query, underscoring the practical importance of accounting for sequence subtype diversity in genomic surveillance efforts. Phylogenetic clustering of PJI isolates revealed that the JH1 *sdrD* subtype was the predominant variant in this clinical context, with JH1-based queries recovering the greatest number of hits among PJI isolates. These findings suggest that the TCH60 sequence represents only a subtype within the broader *S. aureus* population, which is particularly notable, as TCH60 served as the basis for the only previously published SdrD structural characterization (29); therefore, functional or structural inferences drawn solely from TCH60-based studies may not be generalizable. The predominance of the JH1/JH9 lineage among PJI isolates provided a rationale for prioritizing JH1 SdrD for structural determination, ensuring that structural findings are grounded in the clinically dominant sequence variant.

The widely accepted DLL peptide-binding mechanism was proposed based on the N2-N3 structures of SdrG from *S. epidermidis* (32, 33), which lacks a B domain. Conservation of the N2–N3 fold—both at the level of individual domains and their overall bidomain arrangement therefore motivated extension of the DLL model to other Sdr proteins, often without explicitly accounting for the presence and potential functional roles of the B domain. Several issues related to this DLL mechanism remain unanswered. The first relates to the ligand peptide docking in the N2–N3 binding groove. The peptide recognition via the DLL mechanism typically requires the ligand to be presented as an accessible, extended peptide that can dock in the N2–N3 groove. However, some ligand peptides reported for Sdr/Sdr-like adhesins correspond to sequences buried within folded extracellular domains of host receptors, implying that additional events (e.g., local unfolding, mechanical strain, proteolytic processing) may be required to render these motifs accessible under physiological conditions. For example, SdrE has been reported to bind complement factor H (CFH), with the mapped target sequence residing within the C-terminal CFH20 (CCP20) globular module (34, 68). This raises an accessibility problem because the corresponding segment is expected to be structured in the intact CCP20 fold. Thus, productive engagement by SdrE may require local conformational rearrangement or partial unfolding of CCP20—potentially promoted by mechanical strain, receptor dynamics, or proteolytic processing—rather than assuming complete domain denaturation. Similarly, Bpb binds a peptide located within a β-hairpin in the C-terminal subdomain of the fibrinogen α-chain (35, 69). Access to this motif likely entails disruption of the β-hairpin and/or local opening of the subdomain (70). Together, this implies a model in which at least some Sdr/Sdr-like adhesins capture cryptic sequences that become exposed only under specific physiological or mechanical conditions. Further investigation is warranted to determine how cryptic ligand motifs become accessible under physiological conditions. Another unresolved issue is that the DLL peptide-binding groove in N2–N3 appears to impose limited primary-sequence constraints (32, 34, 35), consistent with a binding mode driven largely by backbone interactions. This raises the possibility of broader ligand tolerance, with physiological specificity potentially arising from peptide presentation and accessibility rather than sequence alone.

In addition, the *locking* step—in which a segment after the N3 domain crosses over the bound ligand peptide—may impose geometric and dynamic constraints. Formation of this “lock/latch” element would seemingly require sufficient relative mobility and proximity between membrane-anchored SdrD and its host cell-surface receptor, even though both are often thought to be comparatively constrained on their respective surfaces (71, 72). Moreover, the *latching* step requires a sufficiently long and conformationally flexible segment C-terminal to N3 to traverse the binding groove and stabilize the docked ligand, potentially imposing other geometric and dynamic constraints for surface-anchored adhesins. Importantly, the presence of B domain(s) is likely to have additional major mechanistic consequences – by altering the spacing, orientation, and flexibility of the N2–N3 unit and the post-N3 latch segment, B domains could modulate (or even preclude) canonical DLL locking/latching and enable alternative binding modes.

The experimental observations presented here indicate that the canonical DLL mechanism may have important limitations for SdrD. First, the data shows that the N2–N3–B1 module forms a stable three-domain assembly in solution. Peptide docking at the N2–N3 groove could in principle occur as in other Sdr proteins; however, completion of the *locking* step would likely require substantial rearrangement in which the B1 domain moves away from N2–N3 to create space between N2 and B1 for the ligand peptide and/or its extension. In this model, the N3–B1 interdomain linker would then cross over the bound peptide to “lock” it in place. A subsequent *latching* step would further require that the same N3–B1 linker insert into the N2 latch-binding groove. This places a strong geometric constraint on linker length. Based on structure-guided sequence alignment (**Figure S12**), forming a canonical, full-length latch from the N3–B1 linker appears challenging for SdrD because the linker is short. Indeed, a full latch in SdrD would leave only a one-residue gap between N2 and B1, implying severe steric crowding. A canonical latch seems even less feasible for UafA (36), which has an even shorter N3–B1 loop. Nevertheless, the possibility that SdrD forms a *non-canonical* latch, such as a partial latch or an alternative stabilizing interaction that fulfills the role of locking/latching without reproducing the classical DLL topology, cannot be excluded.

It is generally accepted that the N2–N3 domains constitute the primary receptor-binding region, whereas B domains mainly support surface presentation of N2–N3. However, B domains are not a conserved feature of the Sdr subfamily. ClfA and ClfB, for example, lack B domains altogether. Moreover, among B-domain–containing Sdr proteins, B domains are not equivalent, even if their overall folds are similar. In SdrD and UafA from *S. saprophyticus*, the B domains are predicted from sequence to bind Ca^2+^, a prediction supported by structural data reported here and in prior studies (29, 36). In contrast, the B-domain sequences of SdrC and SdrE do not show obvious Ca^2+^-binding motifs. Notably, both the N and B domains of UafA appear to contribute to binding to erythrocytes (36)—potentially via two distinct mechanisms—although the physiological ligand(s) and the underlying binding mechanism(s) remain to be identified.

Two of the three Ca^2+^-binding sites in each of SdrD B domain are solvent-exposed (**Figure 3B**). The Ca^2+^-mediated N2–B1 interaction (Figure 5A) involves a protruding glutamate (E324 in N2) that contacts Ca^2+^ indirectly via a bridging water molecule. This metal-binding geometry suggests that the B domain could, in principle, engage other proteins bearing similarly protruding acidic residues (Glu or Asp) through an analogous Ca^2+^-dependent interaction. More broadly, the B domain may also participate in ligand recognition via a MIDAS-like mechanism (73, 74). Given the abundance of negatively charged proteins on cell surfaces (75, 76), such Ca^2+^-mediated interactions (potentially multiple valent) could provide an additional mode of receptor engagement beyond N2–N3–centric DLL binding. Considering the cadherin-like nature of DSG-1(77), reported as a host-cell partner adhesin of SdrD (12, 18), another type of potential multivalent Ca^2**+**^-dependent interaction resembling cadherin-cadherin engagement (78–80) between SdrD B domain(s) and DSG1 cadherin domain(s) cannot be excluded.

There are several limitations of the present study. First, while the phylogenetic analysis of the *sdrCDE* locus across a large public genome library and a cohort of 156 PJI isolates provides insight into the population-level distribution of sdrD sequence subtypes, the genomic data are inherently retrospective and subject to the biases of publicly available sequencing repositories, which may not fully capture the global diversity of clinical *S. aureus* isolates. Second, although crystal structures of JH1 SdrD were determined and complemented with SAXS and MD simulations, structural studies were limited to isolated domains and domain combinations in the absence of a physiological ligand. The identity of the physiological ligand for SdrD – and whether DSG-1 or other proposed host receptors engage SdrD through the N2–N3 groove, the B domain, or a combination of both – remains unresolved. Consequently, while the data presented challenge the canonical DLL mechanism as the sole or primary binding mode for SdrD, at this stage, a fully validated alternative mechanism, cannot be presented. Third, the conformational behavior of SdrD observed in solution by SAXS and MD simulations reflects the isolated protein under defined buffer conditions, it may not recapitulate the dynamics of surface-anchored SdrD under the mechanical and biochemical constraints at the host-pathogen interface. Finally, although the structural and biophysical analyses focused on the JH1 lineage as the clinically dominant PJI variant, structural consequences of sequence divergence between the USA300, JH1, and TCH60 subtypes (particularly with respect to ligand-binding surfaces and calcium coordination) remain to be characterized, and functional differences between subtypes cannot be excluded.

Together, the data presented suggest greater sequence, structural, and functional diversity within the Sdr subfamily than previously appreciated. The existence of three distinct *sdrD* sequence subtypes across the *S. aureus* population, combined with structural evidence that SdrD adopts a relatively rigid conformation challenges the canonical ligand-binding model proposed for this protein family, and points to the possibility that different Sdr family members (and different SdrD variants) have evolved distinct mechanisms for engaging host cell-surface receptors. Such diversification may enable the bacterium to broaden its adhesive repertoire across different infection niches, potentially contributing to pathophysiology. The crystal structure of the JH1 SdrD variant presented provides a structural foundation for the design of anti-adhesion therapeutics.

## AUTHOR CONTRIBUTIONS

ME generated expression vectors for all SdrD constructs. PG and AT purified and crystallized SdrD proteins for X-ray crystallography and SAXS studies. KT, PG and YK collected diffraction data, solved and refined structures. XZ and AKN collected and analyzed SEC-SAXS and SAXS data. PG, AT and YK performed thermal shift assays. XZ and AKN performed molecular dynamic simulations. AT and KO purified and submitted *S. aureus* PJI DNA for sequencing and performed alignment and associated downstream analyses of the sequence data. EG performed a literature review on the role of SdrD in PJI and KT analyzed all available structural data. EG drafted the abstract and introduction sections and performed code review of phylogenetic based analyses. KT drafted structural biology section. KGQ analyzed metadata for the PJI isolates. RP and AJ provided supervision and funding. All authors participated in the writing, editing, and review of the manuscript for submission.

## TABLES AND FIGURES & TABLE AND FIGURE LEGENDS

**Figure S1.**
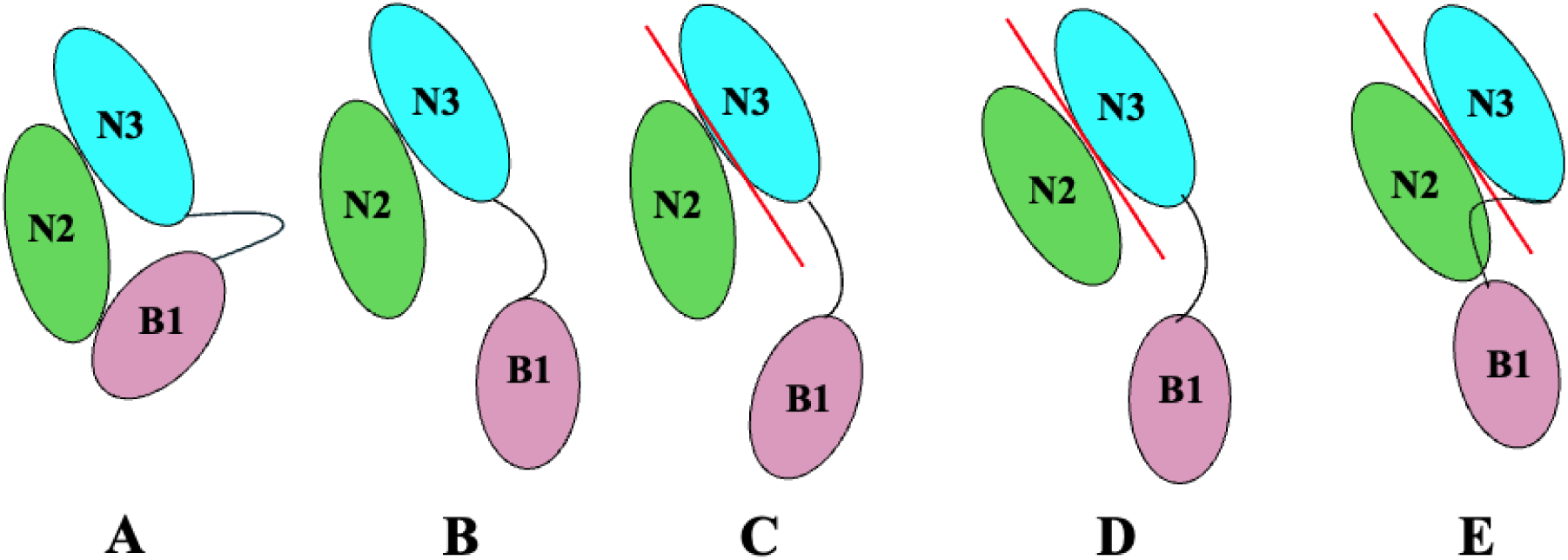
Presumed DLL mechanism for *Staphylococcus aureus* SdrD based on the “dock, lock, and latch (DLL)” ligand-binding mechanism. Only the N2, N3, and B1 domains were used for modeling. (A) The conformation of the N2N3B1 domains observed in crystal structures is regarded an *Apo* state. (B) When the B1 domain detaches from the N2 domain, the conformation is designated an open state. (C) The step in which a peptide segment from a host protein binds in the trench between the N2 and N3 domains, forming an edge β-strand of N3, is referred to as docking. (D) The subsequent locking step represents conformational adjustments within and/or between the N2 and N3 domains that tighten around the bound ligand, stabilizing the complex. (E) The final latching step occurs when the linker peptide between the N3 and B1 domains swings over the bound peptide and inserts into a complementary groove on the N2 domain, forming an additional β-strand. This closes the binding site and yields a highly stable final complex.

**Figure S2.**
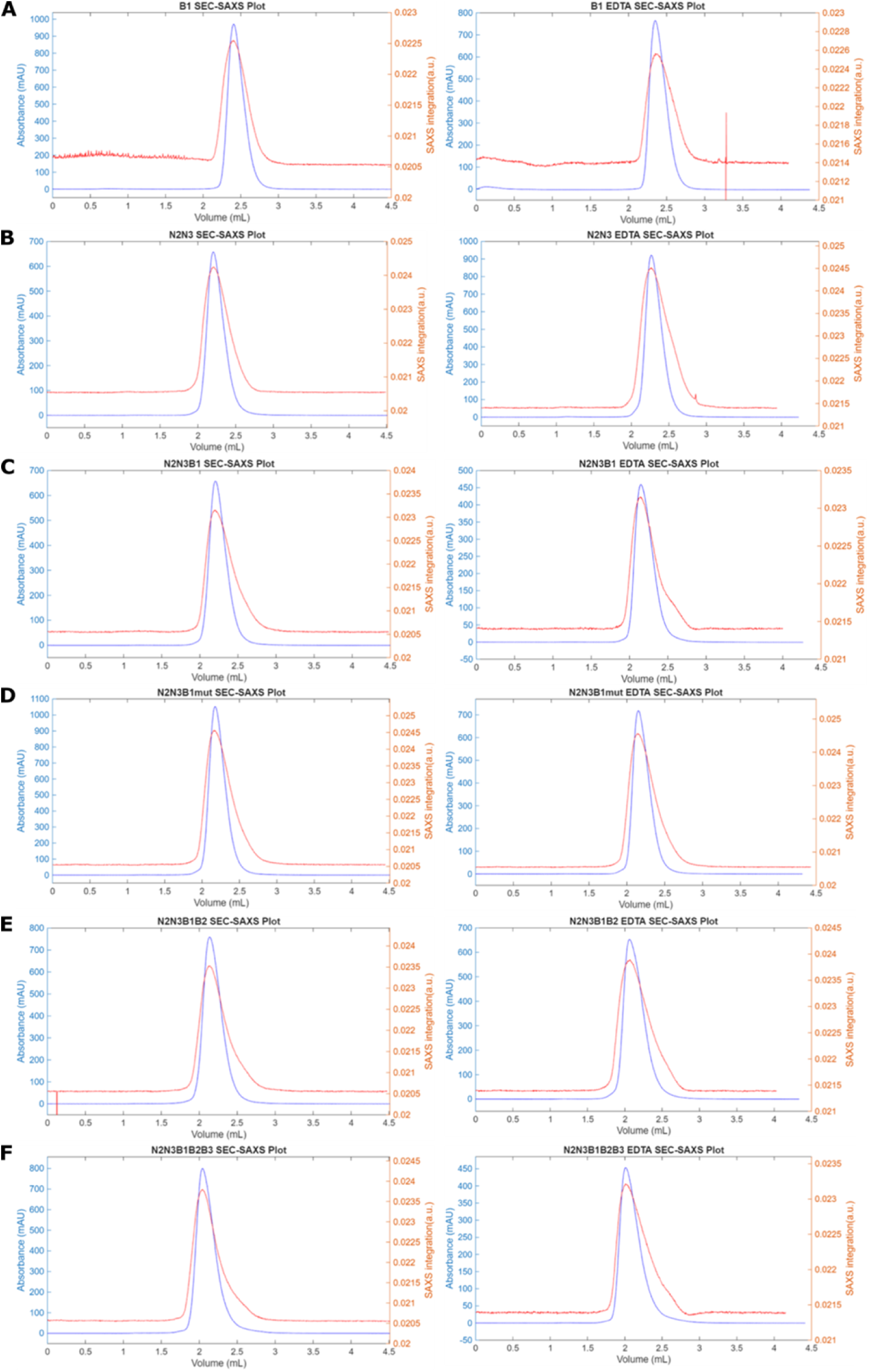
Superimposed SEC profiles detected with UV-vis absorbance at 280 nm (left axis) and SAXS signal integration (right axis) for *Staphylococcus aureus* SdrD. (A) B1, (B) N2N3, (C) N2N3B1, (D) N2N3B1 N574A mutant, (E) N2N3B1B2, (F) N2N3B1B2B3, in protein buffer (20 mM HEPES pH 7.5, 150 mM NaCl, 1 mM TCEP) (left) and in protein buffer with 10 mM EDTA added (right). Profiles detected with SAXS are generally broader than those with UV because SAXS detection is located downstream where the chamber is larger.

**Figure S3.**
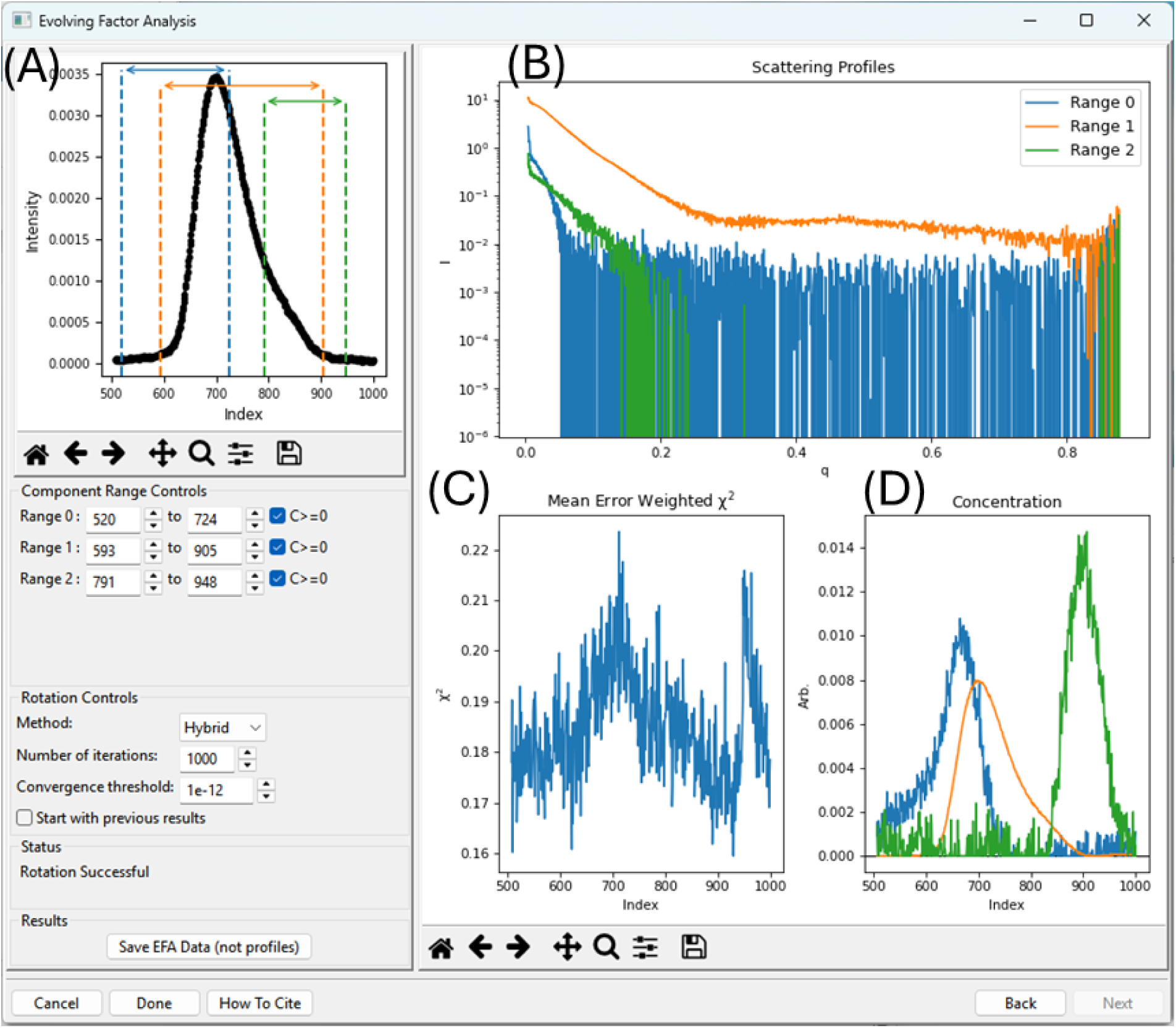
Representative EFA (evolving factor analysis) SEC-SAXS data analysis in BioXTAS RAW. Analysis screenshot captured from RAW for *Staphylococcus aureus* SdrD N2N3B1B2 in protein buffer. (A) Three possible components were identified and their respective existing ranges. (B) The SAXS profiles recovered from EFA analysis for the three components. The second component colored in orange is the major component. (C) Individual fit-goodness plot for SAXS data. (D) Concentration profiles for each component based on SAXS spectra in (B).

**Figure S4.**
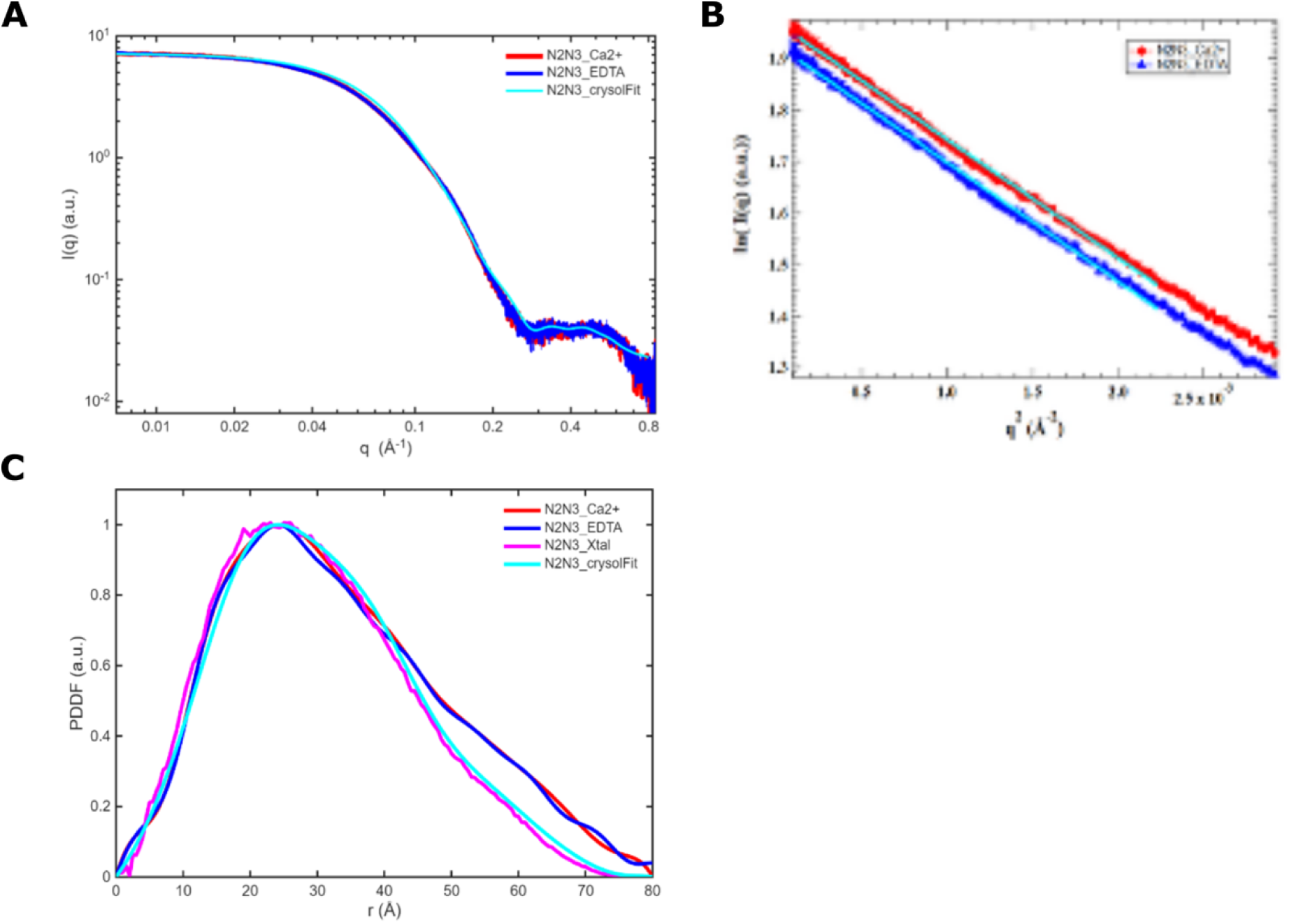
SAXS results for *Staphylococcus aureus* SdrD N2N3 domains. (A) Superimposed SAXS profiles of N2N3 in protein buffer (red curve), and in protein buffer with 10 mM EDTA added (blue). (B) Guinier plots for protein N2N3 in the presence of Ca^2+^ (red) and EDTA (blue), respectively. Cyan lines are Guinier fittings yielding radius of gyration (R_g_) values: 26.2±0.1 Å with Ca^2+^; 26.3±0.2 Å with EDTA. (C) PDDF profiles for N2N3. Red, calculated from experimental SAXS data (red curve in Figure S4A) using GNOM for N2N3 with Ca^2+^; blue, calculated from SAXS data (blue curve inf Figure 2D) for sample with EDTA. Cyan, calculated from fitting SAXS curve (cyan in S4A) using GNOM. Magenta, PDDF calculated from the crystal structure using program SolX2. The difference between cyan and magenta PDDFs arises from the hydration layer added in the CRYSOL fitting.

**Figure S5.**
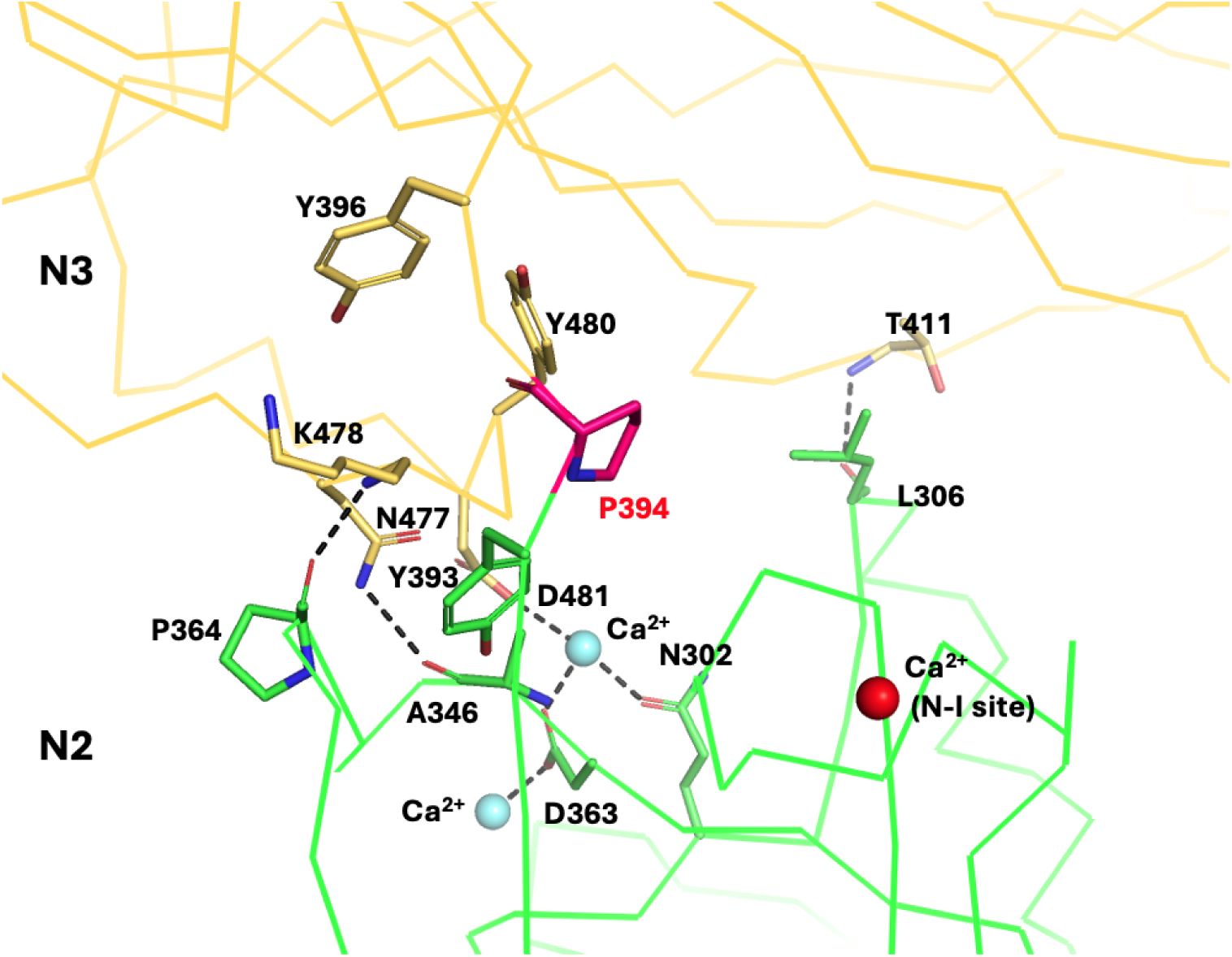
Alpha-trace diagram of the interface between *Staphylococcus aureus* SdrD N2 (green) and N3 (light orange) domains and the primary interactions across their interface. Residues involved in the interactions are shown in stick format. Link residues between the two domains, P394, is highlighted in magenta. Ca^2+^ at the N-I site is shown as a red sphere, while the two Ca^2+^ ions located at N-III site are shown as cyan spheres, see Figure 2A-C. This diagram was prepared based on the *S. aureus* SdrD N2N3 structure.

**Figure S6.**
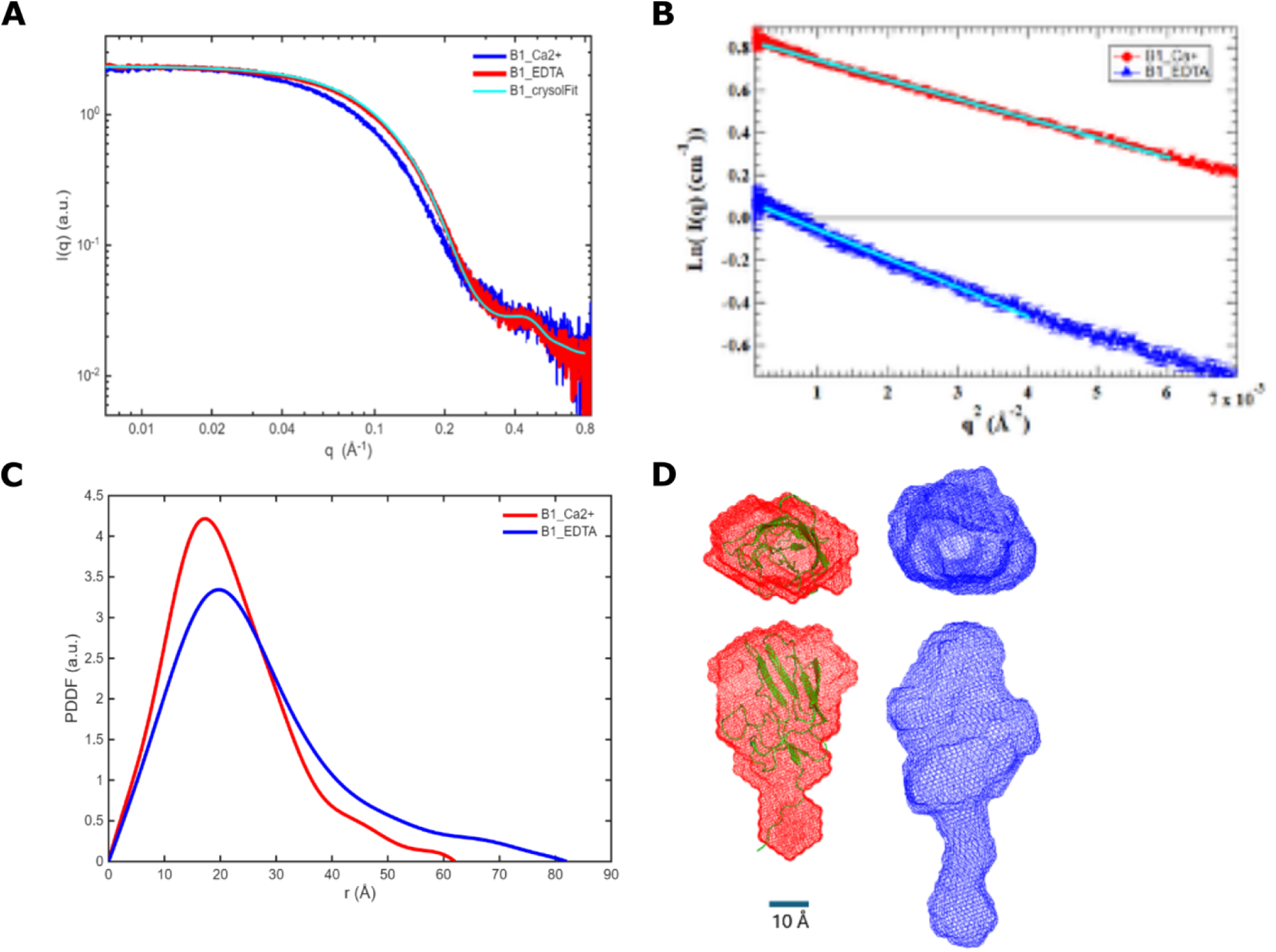
SAXS results for *Staphylococcus aureus* SdrD B1 domain. (A) Superimposed SAXS profiles of B1 in protein buffer (red curve), in protein buffer with 10 mM EDTA added (blue), and fitting with the crystal structure (cyan). The goodness-of-fit (chi2) against the red curve is 0.58. (B) Guinier plots for protein B1 in the presence of Ca^2+^ (red) and EDTA (blue), respectively. Cyan lines are the fitting. Radius of gyration (R_g_) values: 16.6±0.1 Å with Ca^2+^; 20.2±0.2 Å with EDTA. (C) PDDF profiles for N2N3. Red, calculated from experimental SAXS data (red curve in Figure S5A) using GNOM; blue, calculated from blue SAXS data in Figure S5A. (D) Top and side views of the most-probable SAXS molecular envelopes for B1 reconstructed from SAXS data at the two conditions. Color coded the same as in A-C. The crystal structure was displayed in green color and in cartoon mode.

**Figure S7.**
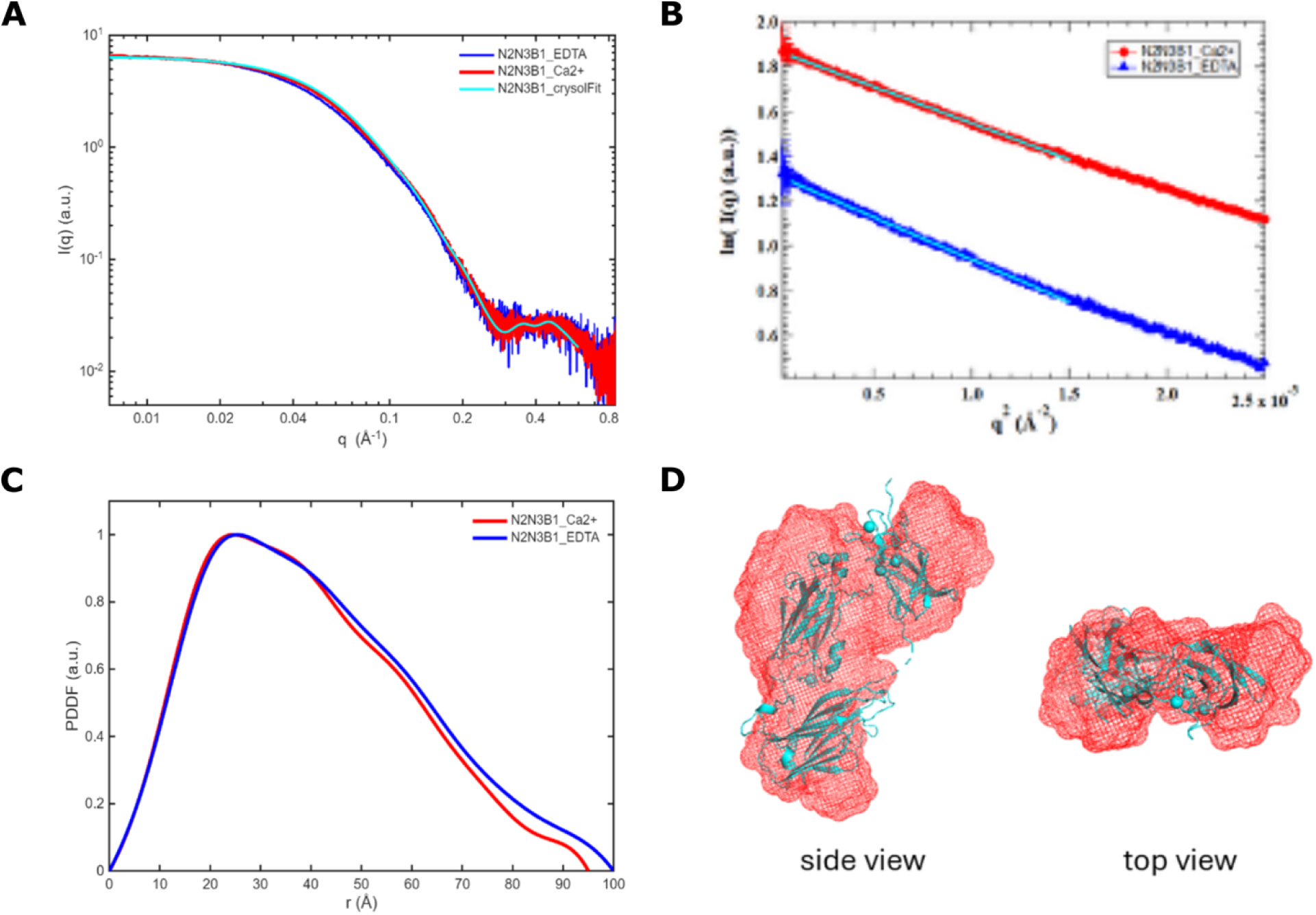
MD simulations on *Staphylococcus aureus* SdrD B1 domain. (**A**,**B**) Comparison of the conformational distribution of the *R_g_* sampled for the B1 domain with and without Ca^2+^ at 300 K (**A**) and 450 K (**B**). (**C**, **D**) Comparison of the conformational distribution of root mean square deviation (RMSD) of the α carbons from the crystal structure at 300 K (**C**) and 450 K (**D**). (**E**, **F**) Representative trajectories from sample MD simulations conducted at 300 K (**E**) and 450 K (**F**). (**G**, **H**) Comparison of theoretical SAXS for the B1 domain with (**G**) and without (**H**) Ca^2+^ to the experimental SAXS data. The average MD structure represents the weighted average for the conformational distributions depicted in Figure 4C. Minor discrepancies are attributed to effects of the solvation layer and the experimental SAXS representing an ensemble average over different conformations.

**Figure S8.**
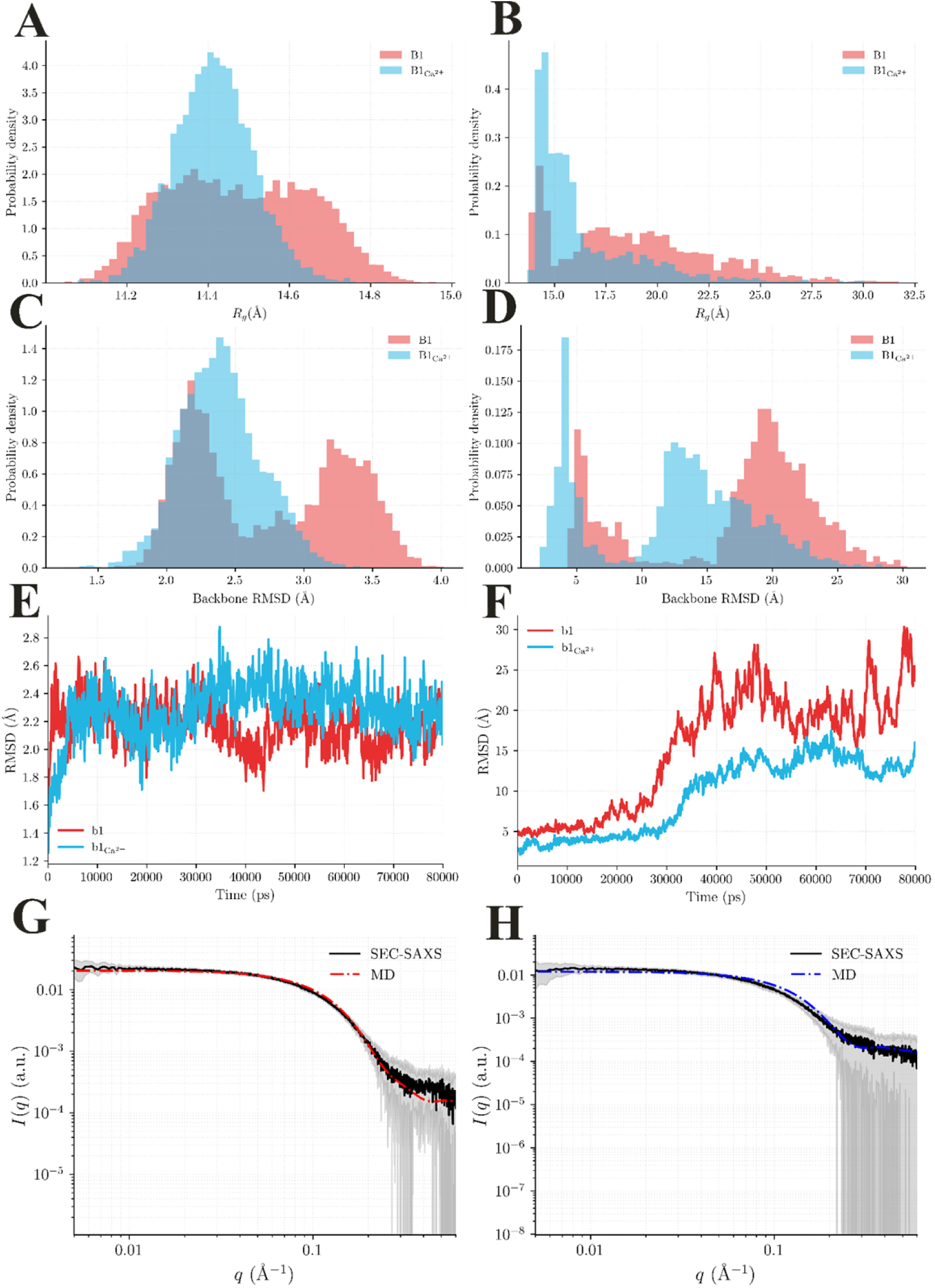
SAXS results for *Staphylococcus aureus* SdrD N2N3B1. (A) Superimposed SAXS profiles of N2N3B1 in protein buffer (red curve), in protein buffer with 10 mM EDTA added (blue), and fitting with the crystal structure (cyan). The goodness-of-fit (chi2) of CRYSOL fitting with the crystal structure against the red curve is 2.2. (B) Guinier plots for protein N2N3B1in the presence of Ca^2+^ (red) and EDTA (blue), respectively. Cyan lines are fittings which yield the following radius of gyration (R_g_) values: 31.2±0.2 Å with Ca^2+^; 33.6±0.2 Å with EDTA. (C) PDDF profiles for N2N3B1. Red, calculated from experimental SAXS data (red curve in Figure S7A) using GNOM; blue, calculated from blue SAXS data in Figure S7A. (D) Side and top views of the most-probable SAXS molecular envelope for N2N3B1 reconstructed for the data collected with Ca^2+^. Color coded the same as in A-C. The N2N3B1 crystal structure was displayed in cyan color and in cartoon mode.

**Figure S9.**
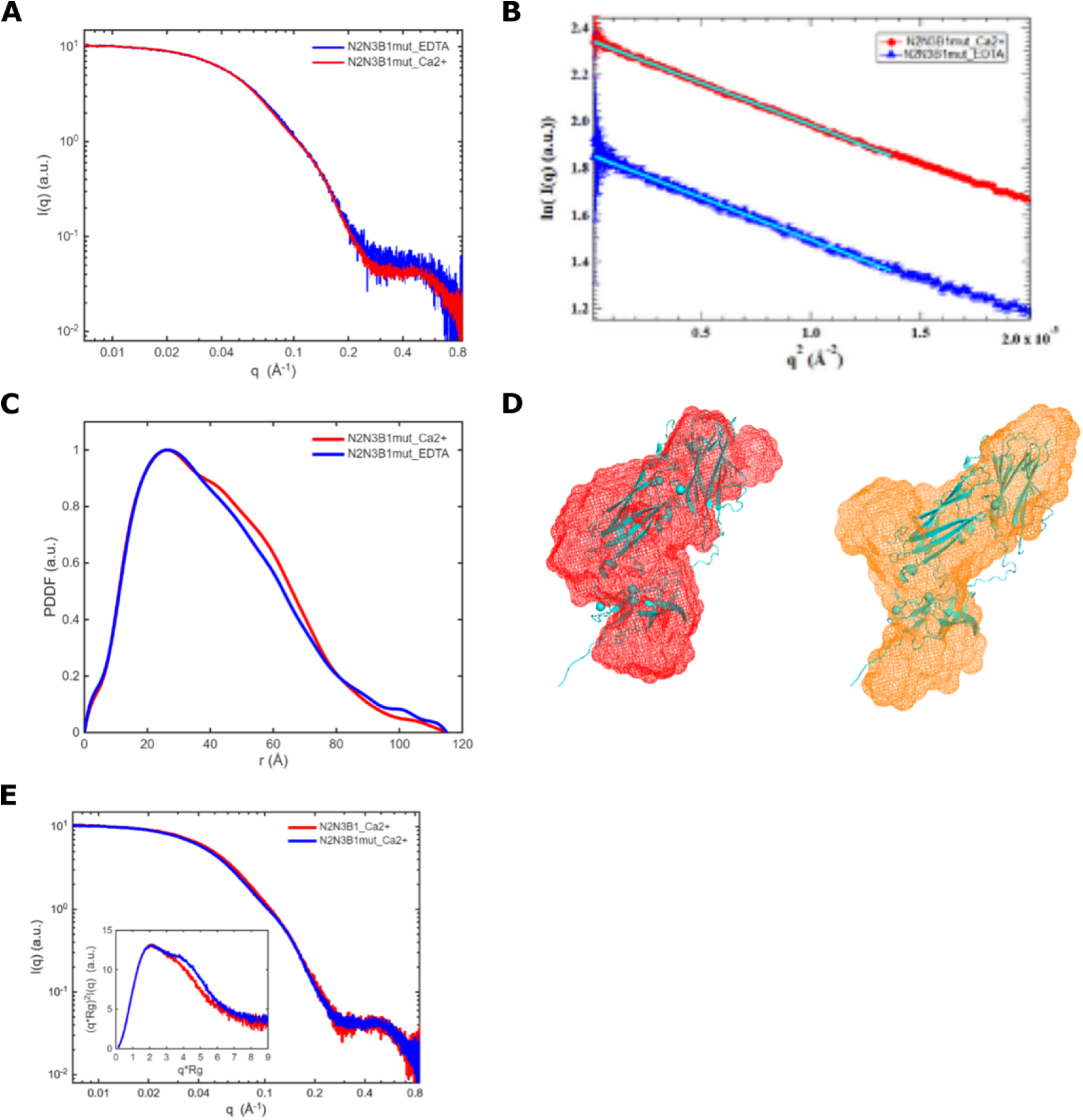
SAXS results for *Staphylococcus aureus* SdrD N2N3B1mut. (A) Superimposed SAXS profiles of N2N3B1mut in protein buffer (red curve), and in protein buffer with 10 mM EDTA added (blue). (B) Guinier plots for protein N2N3B1in the presence of Ca^2+^ (red) and EDTA (blue), respectively. Cyan lines are the fittings which yield radius of gyration (R_g_) values: 32.8±0.2 Å with Ca^2+^; 32.9±0.3 Å with EDTA. (C) PDDF profiles for N2N3B1mut. Red, calculated from experimental SAXS data (red curve in Figure S8A) using GNOM; blue, calculated from blue SAXS data in Figure S8A. (D) Side views of the refined SAXS molecular envelope for N2N3B1 (red) and N2N3B1mut (orange) reconstructed for data collected with Ca^2+^. The N2N3B1 crystal structure is displayed in cyan color and in cartoon mode.

**Figure S10.**
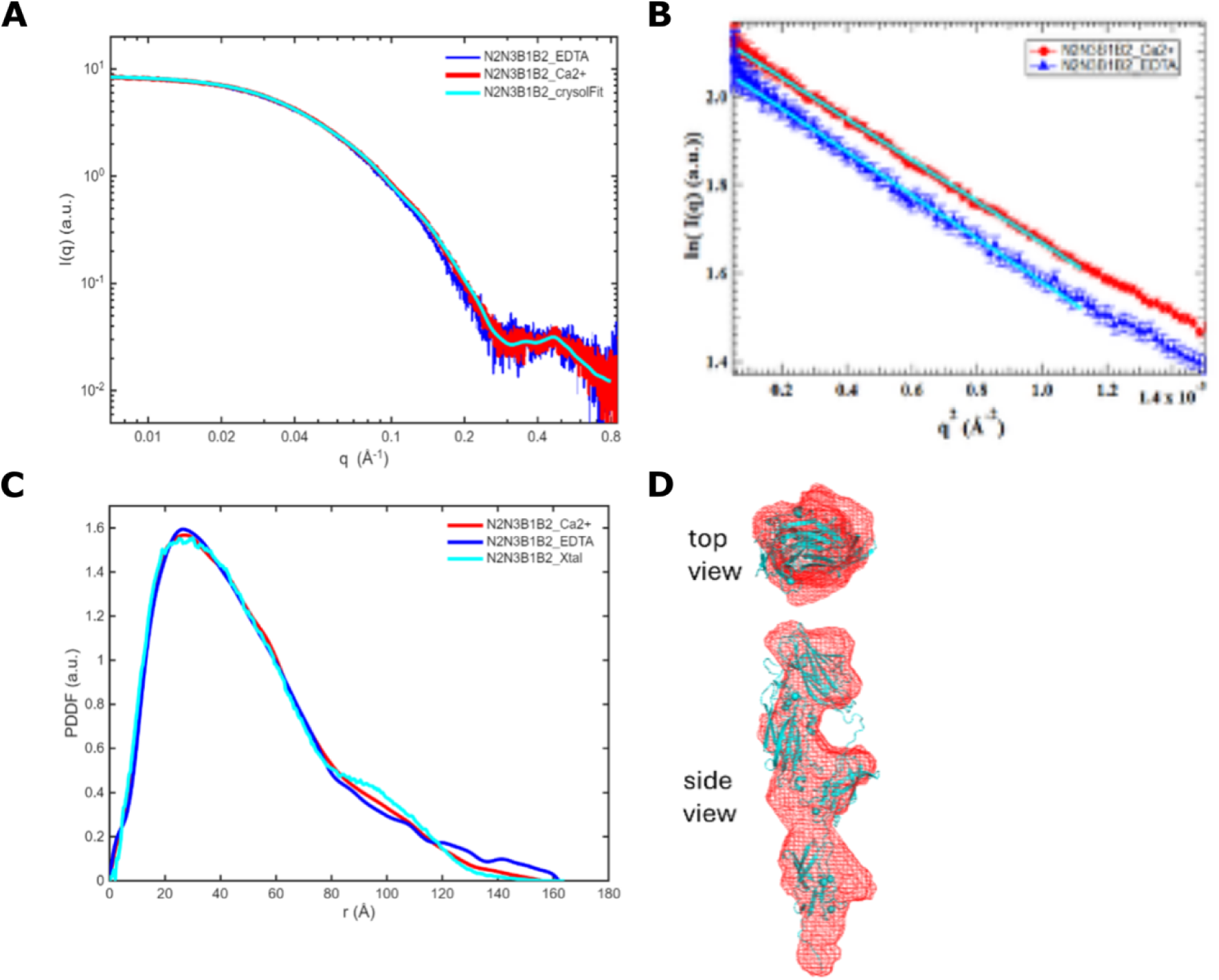
SAXS results for *Staphylococcus aureus* SdrD N2N3B1B2. (A) Superimposed SAXS profiles of N2N3B1B2 in protein buffer (red curve), in protein buffer with 10 mM EDTA added (blue), and fitting with the crystal structure (cyan). The goodness-of-fit (chi2) of CRYSOL fitting with the crystal structure against the red curve is 0.44. (B) Guinier plots for protein N2N3B1B2 in the presence of Ca^2+^ (red) and EDTA (blue), respectively. Cyan lines are fittings which yield the following radius of gyration (Rg) values: 37.5±0.2 Å with Ca2+; 38.3±0.3 Å with EDTA. (C) PDDF profiles for N2N3B1B2. Red, calculated from experimental SAXS data (red curve in Figure S9A) using GNOM; blue, calculated from blue SAXS data in Figure S9A. (D) Side and top views of the most-probable SAXS molecular envelope for N2N3B1B2 reconstructed for the data collected with Ca^2+^. The N2N3B1B2 crystal structure is displayed in cyan color and in cartoon mode and superimposed with SAXS structural envelope.

**Figure S11.**
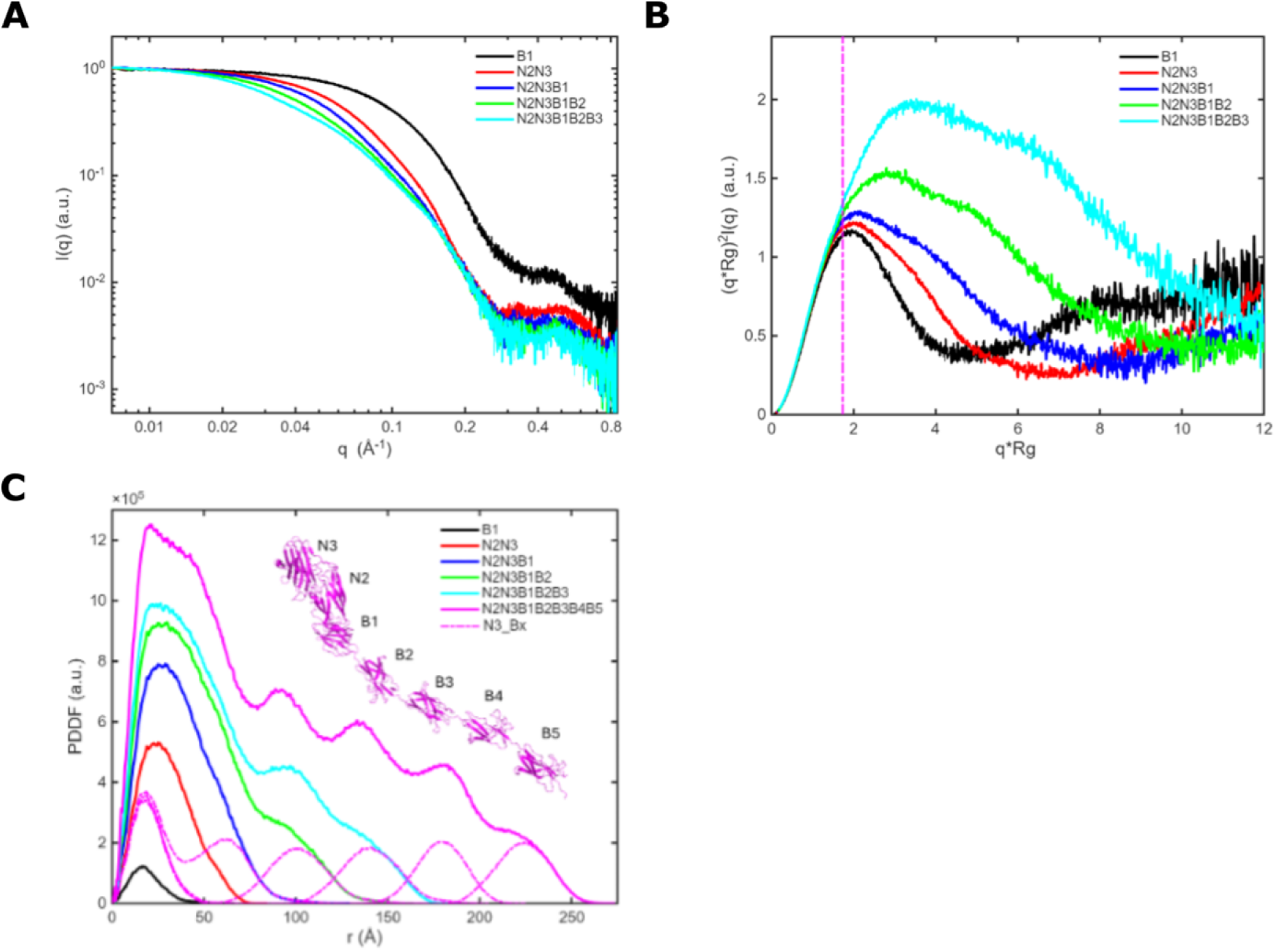
(A) Summary plot of normalized experimental SAXS profiles for all constructs. (B) Dimensionless Kratky plots for experimental SAXS data in (A). Colors are coded as in (A). The magenta vertical line is 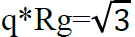, where the peak of spherical particle appears. The higher the aspect ratio of length to cross-section, the further the peak position is away from 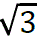. (C) Theoretical PDDFs of all constructs calculated from atomic structures. Among the atomic structures, B1, N2N3, N2N3B1 and N2N3B1B2 are crystal structures, and N2N3B1B2B3 and N2N3B1B2B3B4B5 are AF3 models. N2N3B1B2B3B4B5 is displayed in the inset with labels of N and B domains. The dash line profiles in magenta are five PDDFs for the ensemble structure, consisting of N3 and one individual B domains, i.e., N3_B1, N3_B2, N3_B3, N3_B4, and N3_B5, respectively. They all consist of two peaks, one short distance peak centering at ∼18 Å, and one far distance peak centering from 62 to 224 Å, depending on up the location of B domain. The first peak reflects the intra-domain pair-wise distance within N3 or B domains, while the second peak describes the inter-domain pair-wise distance correlations between N3 and B domains. The peak position of the second peak reflects the inter-domain center-to-center distance.

**Figure S12.**
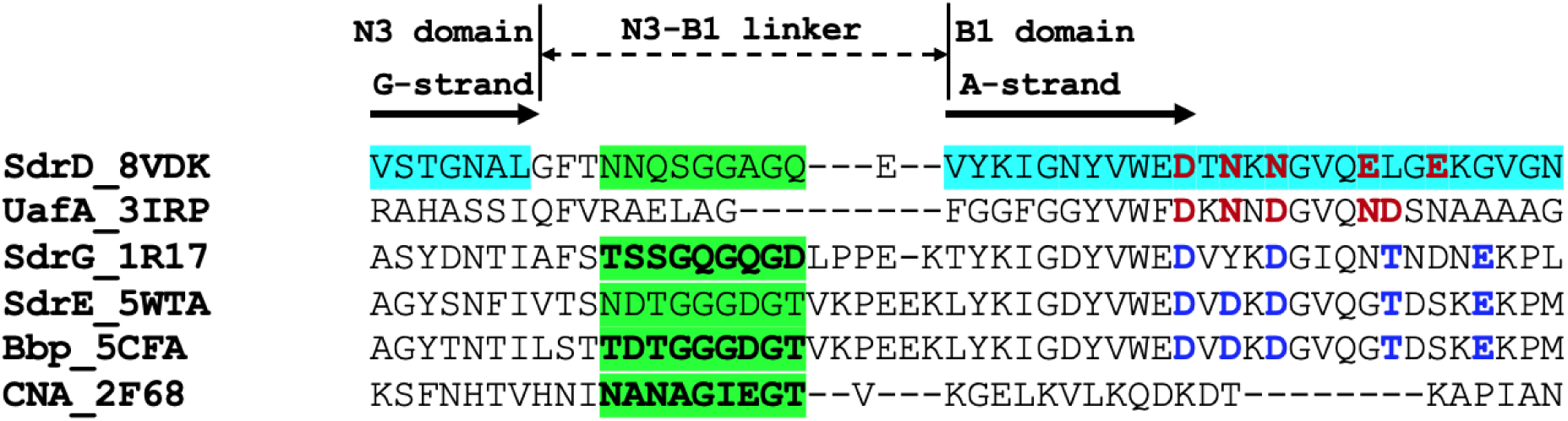
Structure based sequence alignment of N3-B1 interdomain region. Relevant Sdr and Sdr-like molecules were used for the alignment. Crystal structure PDB codes are shown after molecular names for reference. The last strand of the N3 G-strand and a part of N-terminal region of B1 domain (including A-strand) were used in the alignment. Strand assignments are based on SdrD structure. Observed (bolded) and presumed latch forming motifs within the N3-B1 linkers are highlighted in green. In the B1 domain region, observed metal ion-binding residues are in red font in SdrD and UafA. Residues in equivalent or nearly equivalent positions of metal-binding sites in SdrG, SdrE and Bbp are in blue font.

